# Functional and structural insights into the multi-step activation and catalytic mechanism of bacterial ExoY nucleotidyl cyclase toxins bound to actin-profilin

**DOI:** 10.1101/2023.01.13.523968

**Authors:** Magda Teixeira-Nunes, Pascal Retailleau, Dorothée Raoux-Barbot, Martine Comisso, Anani Amegan Missinou, Christophe Velours, Stéphane Plancqueel, Daniel Ladant, Undine Mechold, Louis Renault

## Abstract

ExoY virulence factors are members of a family of bacterial nucleotidyl cyclases (NCs) that are activated by specific eukaryotic cofactors and overproduce cyclic purine and pyrimidine nucleotides in host cells. ExoYs are actin-activated NC toxins. Here, we investigate the *Vibrio nigripulchritudo* Multifunctional-Autoprocessing Repeats-in-ToXin (MARTX) ExoY effector domain (Vn-ExoY) as a model for ExoY-type members that interact with monomeric (G-actin) rather than filamentous (F-actin) actin. Vn-ExoY binds with only modest affinity to free or profilin-bound G-actin, but can capture the G-actin:profilin complex for its own activation by preventing the spontaneous or VASP- or formin-mediated assembly of G-actin:profilin at the barbed ends of F-actin *in vitro*. This may prolong the lifetime of the cofactor-bound state of Vn-ExoY at sites of active actin cytoskeleton remodelling. A series of high-resolution crystal structures of nucleotide-free, 3’-deoxy-ATP- or 3’-deoxy-CTP-bound Vn-ExoY, activated by free or profilin-bound G-actin-ATP/-ADP show that the cofactor only partially stabilises the nucleotide-binding pocket (NBP) of all NC toxins. Substrate binding promotes a large, previously-unidentified, closure of their NBP. This confines catalytically important residues of the NC toxins and metal cofactors around the substrate and promotes the recruitment of two metal ions to tightly coordinate the triphosphate moiety of purine or pyrimidine nucleotide substrates. Residues that play an important role in both the purinyl and pyrimidinyl cyclase activity of NC toxins are validated in Vn-ExoY and the distantly-related ExoY from *Pseudomonas aeruginosa* that interact with F-actin. The data conclusively demonstrate that NC toxins employ a similar two-metal-ion mechanism for catalysing the cyclisation reaction of nucleotides of different sizes. These structural insights into the dynamics of the actin-binding interface of actin-activated ExoYs and the multi-step activation of all NC toxins open up new perspectives for identifying ways to specifically inhibit these bacterial NC enzymes.

**Author Summary:** ExoY toxins belong to a family of bacterial nucleotidyl cyclases (NCs) that are injected into eukaryotic cells and bind to specific host cofactors to trigger their toxic, potent NC enzymatic activity. They alter host cell signalling by overproducing purine and pyrimidine cyclic nucleotides, which act as canonical and non-canonical intracellular messengers, respectively. The molecular and mechanistic details underlying the activation and catalytic specificities of NC toxins are only partially understood. Here, we investigate ExoY-type members that are unable to interact with actin filaments for their activation. We show *in vitro* that such ExoYs capture the actin:profilin complex for activation by disrupting its association with the most dynamic ends of actin filaments. We have captured several structural snapshots along the Vn-ExoY activation pathway by G-actin or G-actin-profilin without or with purine or pyrimidine nucleotide analogues. Our structural data reveal unprecedented mechanistic details of how the active site of all NC toxins is sequentially remodelled by cofactor and substrate binding, how they can accommodate nucleotides of different sizes as substrates, and elucidate important features of their catalytic reaction. These structural insights into the multi-step activation of NC toxins provide new perspectives for identifying ways to specifically inhibit this class of NC enzymes.

## Introduction

ExoY-like nucleotidyl cyclase (NC) toxins are virulence factors produced by several Gram-negative β- and γ-Proteobacteria [1]. They belong to a family of bacterial NCs that share a structurally-related catalytic NC domain that is inactive in bacteria. When injected into eukaryotic cells, they bind to specific eukaryotic proteins that stimulate their enzymatic activity by several orders of magnitude [2, 3]. They then become highly active cyclases with broad substrate specificity [4, 5] and increase the intracellular concentrations of cyclic purine (cAMP, cGMP) and pyrimidine (cCMP, cUMP) nucleotides, thereby interfering with host cell signalling (reviewed in [1, 2, 6–8]). While the *Bacillus anthracis* edema factor (EF) and *Bordetella pertussis* CyaA toxins are both activated by calmodulin (CaM), the ExoY-like NC toxins use actin [3]. CaM-activated toxins have been extensively studied over the last decades. On the other hand, the functional characteristics and virulence mechanisms of the ExoY-like NC toxins still need to be clarified.

Exoenzyme Y (Pa-ExoY) is the second most prevalent type III secretion system (T3SS) exotoxin encoded in the genomes of clinical or environmental isolates of *P. aeruginosa* [9]. Pa-ExoY NC activity in host cells alters the host immune response [10, 11] and leads to the release of cytotoxic amyloids from the pulmonary endothelium, impairing cellular repair after infection [7, 12, 13]. In *Vibrio vulnificus* biotype 3, a pathogen that causes severe foodborne and wound infections in humans, the ExoY module of its MARTX toxin (Vv-ExoY) is essential for virulence [14]. ExoY NC homologues from different pathogens can potentially exert different cytotoxic effects in infected cells due to differences in their protein sequence, the form of actin they require to become active, and their substrate specificities [5]. The MARTX Vn-ExoY module of *Vibrio nigripulchritudo*, an emerging pathogen in marine shrimp farming, is representative of the ExoY modules found in *Vibrio* strains. It shares only 38 % sequence similarity with Pa-ExoY. Whereas Pa-ExoY requires actin filaments (F-actin) for maximal activation [3], Vn-ExoY is selectively activated by G-actin *in vitro*. Vn-ExoY, like EF and CyaA [15], strongly prefers ATP and can use CTP as substrate, but much less efficiently [5]. Pa-ExoY, on the other hand, synthesises a wide range of cyclic nucleotide monophosphates (cNMPs) [15] with the following *in vitro* substrate preference: GTP>ATP≥UTP>CTP [5]. Recently, cUMP and cCMP have come to light as novel second messengers that activate antiviral immunity in bacteria [16]. Their role in eukaryotic cells, on the other hand, is still largely to be elucidated [17].

Bacterial NC toxins are defined as class II adenylate cyclases (ACs). Their catalytic domain is structurally unrelated to the widely distributed class III adenylyl or guanylyl cyclases (GCs) [18–25] and may represent a potential drug target against bacterial infections. In their cofactor-bound state, they are several orders of magnitude more active than activated mammalian class III ACs [2], which use a dimeric catalytic architecture. The molecular basis for the turnover differences between these two AC classes remains to be elucidated. The catalytic domain of NC toxins can be structurally divided into subdomains C_A_ and C_B,_ with the catalytic site located in a groove at their interface [23]. EF and CyaA recognise CaM via two flexible regions of C_A_ that are called switch A (residues 502-551 in EF, 199-274 in CyaA) and switch C (630-659 in EF, 348-364 in CyaA) [23, 24, 26]. In CaM-bound EF, switch A and C stabilise another key region of C_A_, called switch B (578-591 in EF, 299-311 in CyaA). This region contains residues that either directly bind ATP or stabilise catalytic residues in the groove between C_A_ and C_B_ [23]. When EF is inactive, switch B is disordered because switch A and C are distant from the catalytic site. Activation of EF or CyaA by CaM does not appear to require a large subdomain movement between C_A_ and C_B_, but rather a CaM-induced movement of switch A and C towards the catalytic site [2]. This in turn stabilises switch B and ATP binding to favour a catalytically competent form of the enzyme. This allosteric activation-by-stabilisation mechanism was recently endorsed for the actin-activated NC toxins. In the first structure of Pa-ExoY crystallised alone after limited in-situ proteolysis, the three switches were not visible in the electron density due to partial proteolysis and/or flexibility [25]. Recent cryo-EM structures of actin-activated Pa-ExoY and Vv-ExoY revealed that Pa- and Vv-ExoY also use their central switch A and C-terminal switch C to recognise F- and G-actin subunits, respectively [27]. Molecular dynamics simulations suggested that binding of Pa-ExoY to F-actin stabilises its otherwise flexible switches A and C, thereby stabilising the entire toxin and nucleotide-binding pocket (NBP). Despite these advances, the number of ions required for maximum efficiency, the nature of the catalytic base, or the coordination and role of the residues involved in the various reaction steps are still a matter of debate [1, 23, 24, 26–32]. The molecular basis of the NC toxin’s pyrimidinyl cyclase activity is also unknown.

Using *in vitro* functional and structural studies, we show that Vn-ExoY capture the actin:profilin complex for activation by disrupting its association with the most dynamic ends (barbed-ends) of actin filaments. We present a series of high-resolution structures of Vn-ExoY activated by free or profilin-bound actin-ATP/ADP, in the absence or presence of non-cyclisable ATP or CTP analogues bound to Mg^2+^ or Mn^2+^ ions. These structural insights highlight important features of the interface and functional dynamics of ExoY NC toxins with G/F-actin, and explain how NC toxins can use large purine or small pyrimidine nucleotides (with double- or single-carbon nitrogen ring bases, respectively) as substrates. Residues that play an important role in the purinyl and pyrimidinyl cyclase activity of NC toxins are validated in Vn-ExoY and the distantly-related Pa-ExoY.

## Results

### Vn-ExoY binds to actin monomers with only modest affinity and inhibits actin self-assembly

As a model for ExoY toxins that are selectively activated by G-actin, we used a functional Vn-ExoY^3412-3873^ effector domain/module corresponding to residues 3412 to 3873 of *V. n.* MARTX [5]. We first examined the effect of Vn-ExoY on actin-ATP/-ADP self-assembly. To avoid the indirect effects due to ATP depletion by Vn-ExoY AC activity, we used an inactive double mutant K3528M/K3535I, hereafter referred to as Vn-ExoY^DM^ (corresponding to Pa-ExoY^K81M/K88I^) [5]. Actin polymerisation kinetics were monitored using pyrene-labelled actin, which shows an increase in fluorescence intensity when incorporated into filaments. While inactive Pa-Exo^K81M^ accelerated the rate of actin polymerisation (Fig 1A) [27], Vn-ExoY^DM^ inhibited the polymerisation rate of both G-actin-ATP-Mg and G-actin-ADP-Mg (Figs 1B-C). Thus, Vn-ExoY interacts with G-actin-ATP or -ADP in a very different way than Pa-ExoY. To understand the inhibitory effect of Vn-ExoY, we examined the association of free or Vn-ExoY^DM^-bound G-actin with the barbed (+) or pointed (-) ends of filaments. Spectrin-actin or gelsolin-actin seeds were used to induce elongation at the barbed- and pointed-ends, respectively. Interaction of Vn-ExoY^DM^ with G-actin-ATP-Mg inhibited actin assembly at both ends (Fig 1D). The dose-dependent effects of Vn-ExoY^DM^ on barbed- and pointed-end elongation rates were consistent with the formation of a 1:1 molar complex between G-actin and Vn-ExoY^DM^ sequestering G-actin with an equilibrium dissociation constant (Kd) of 21 μM (Fig 1D). To confirm the binding affinity by direct measurement, we performed fluorescence-detected sedimentation velocity experiments using analytical ultracentrifugation (AUC-FDS) with G-actin-ATP labelled with the fluorescent dye Alexa-488 and unlabelled Vn-ExoY fused to Thioredoxin (Trx-Vn-ExoY). This protein fusion, which is more soluble than Vn-ExoY, was preferred for dose-dependent analyses at high concentrations (Fig 1E). With free labelled G-actin (0.06 µM) bound to Latrunculin-A, an actin-polymerisation inhibitor, the actin sedimentation coefficient distribution gave a peak centred at an s_20,w_ value of 3.3 S (Fig 1E, left panel) consistent with a globular actin monomer of 42 kDa. Upon incremental addition of Trx-Vn-ExoY, the sedimentation coefficient distribution of labelled actin shifted to higher s-values (from 3.2 up to 4.7 S), demonstrating the interaction of Vn-ExoY with actin. Peak position and shape were concentration-dependent, and the relative proportion of the larger species in the peaks increased with increasing concentrations of Trx-Vn-ExoY. The titration was consistent with the formation of a 1:1 Vn-ExoY:G-actin complex with a Kd of 11.7 ± 0.9 µM (Fig 1E, right panel). Vn-ExoY therefore forms a sequestering but modest affinity complex with G-actin. Its affinity is modest compared to that of P-a-ExoY for F-actin [3] or EF and CyaA for calmodulin [33, 34]. It is also modest compared to that of G-actin binding proteins (G-ABPs), which regulate the G-actin pool in eukaryotic cells [35].

**Fig. 1.**
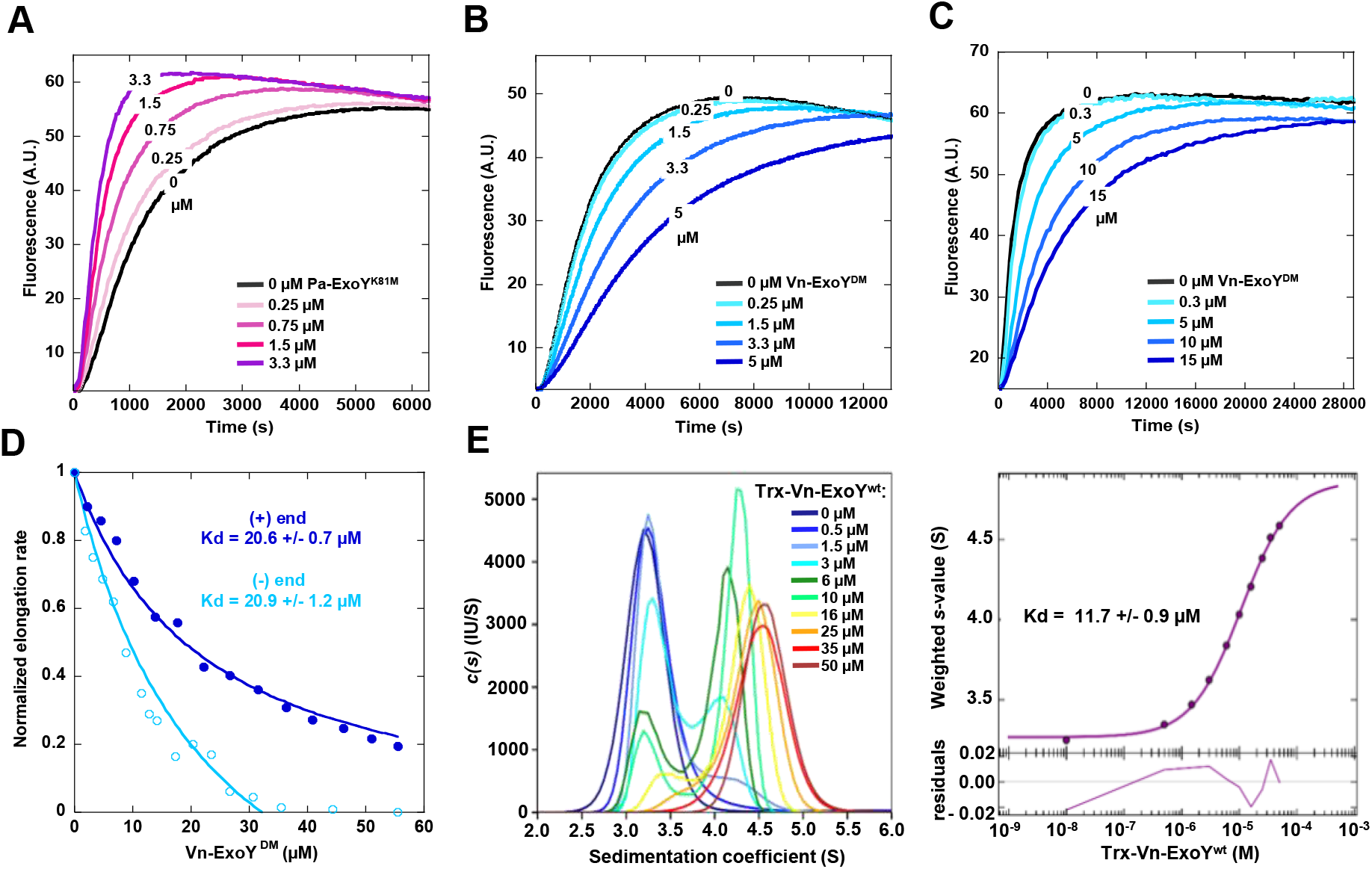
Vn-ExoY sequesters actin-ATP/ADP monomers in actin self-assembly by interacting with modest affinity. **(A-C)** Polymerisation time course of 4 µM ATP-Mg-G-actin (5 % pyrenyl-labelled, (A-B)) or 10 µM ADP-Mg-G-actin (10 % labelled, (C)) in the absence (black) and presence of Pa-ExoY^K81M^ (A) or Vn-ExoY^DM^ (B-C) at the indicated concentrations (µM). **(D)** Initial rates of barbed-end (dark blue) or pointed-end (light blue) growth from spectrin-actin or gelsolin-actin seeds, respectively, were measured with 1.5 μM G-actin (10 % pyrenyl-labelled) and increasing concentrations of Vn-ExoY^DM^. Initial rates were normalised to the initial elongation rate with free G-actin from a linear fit to the first 120 s of each time course. Vn-ExoY^DM^ inhibits both barbed and pointed-end growth by sequestering G-actin with the indicated Kd (µM). **(E)** AUC-FDS measurement of the binding affinity of Vn-ExoY for G-actin. The sedimentation coefficient distribution c(s) (left panel) of Alexa-488-labelled-G-actin-ATP (60 nM) bound to the actin-polymerisation inhibitor Latrunculin A (2.5 µM) was determined in the absence and presence of increasing concentrations of unlabelled Trx-Vn-ExoY (from 0.5 to 50 µM). The resulting sw isotherms from c(s) integration are shown in the right panel. In the isotherm plots, the solid circles are the sw data from the dilution series. The solid line is the best-fit isotherm giving an estimated Kd of 11.7 µM for the interaction between Trx-Vn-ExoY and G-actin-ATP-Latrunculin-A.

### Vn-ExoY is efficiently activated by G-actin bound to profilin

We next examined the *in vitro* AC activity of Vn-ExoY stimulated by G-actin in the presence of three regulatory G-ABPs that display different binding interfaces on G-actin: (i) a chimeric β-thymosin domain of Thymosin-β4 and Ciboulot proteins, which inhibits actin self-assembly by sequestering G-actin [36], (ii) profilin and (iii) a profilin-like WH2 domain of the Cordon-Bleu protein [37]. The latter two control the unidirectional assembly of actin monomers at the barbed-ends of filaments. At high, saturating concentrations of these three G-ABPs, only profilin does not interfere with G-actin-induced activation of Vn-ExoY AC activity, and allows potent AC (Fig 2A). We then performed AUC-FDS experiments, monitoring the sedimentation behaviour of Alexa-488-labelled-G-actin in the presence of Vn-ExoY, profilin or both proteins. The formation of complexes of increasing size was observed by the shifts from an s_20,w_ value of 3.5 S obtained for the peak of free Alexa488-labelled actin to 4.2 S with profilin (15 kDa), 5.0 S with the larger Trx-Vn-ExoY protein (69.1 kDa), and to a higher value of 5.5 S with both proteins (Fig 2B), validating the formation of a ternary Vn-ExoY:actin:profilin complex in solution. These result are consistent with the interaction of *V. vulnificus* Vv-ExoY with actin:profilin [27]. We next investigated whether the interaction of Vn-ExoY with G-actin was modulated by the simultaneous binding of profilin. We measured the binding strength of Trx-Vn-ExoY^DM^ to fluorescently labelled G-actin-ATP alone or bound to profilin using microscale thermophoresis (MST). Titrations of the interaction by measuring changes in fluorescence (Fig 2C) or MST signal (S1 Fig) show that Vn-ExoY binds with similar affinity to free or profilin-bound G-actin.

**Fig. 2.**
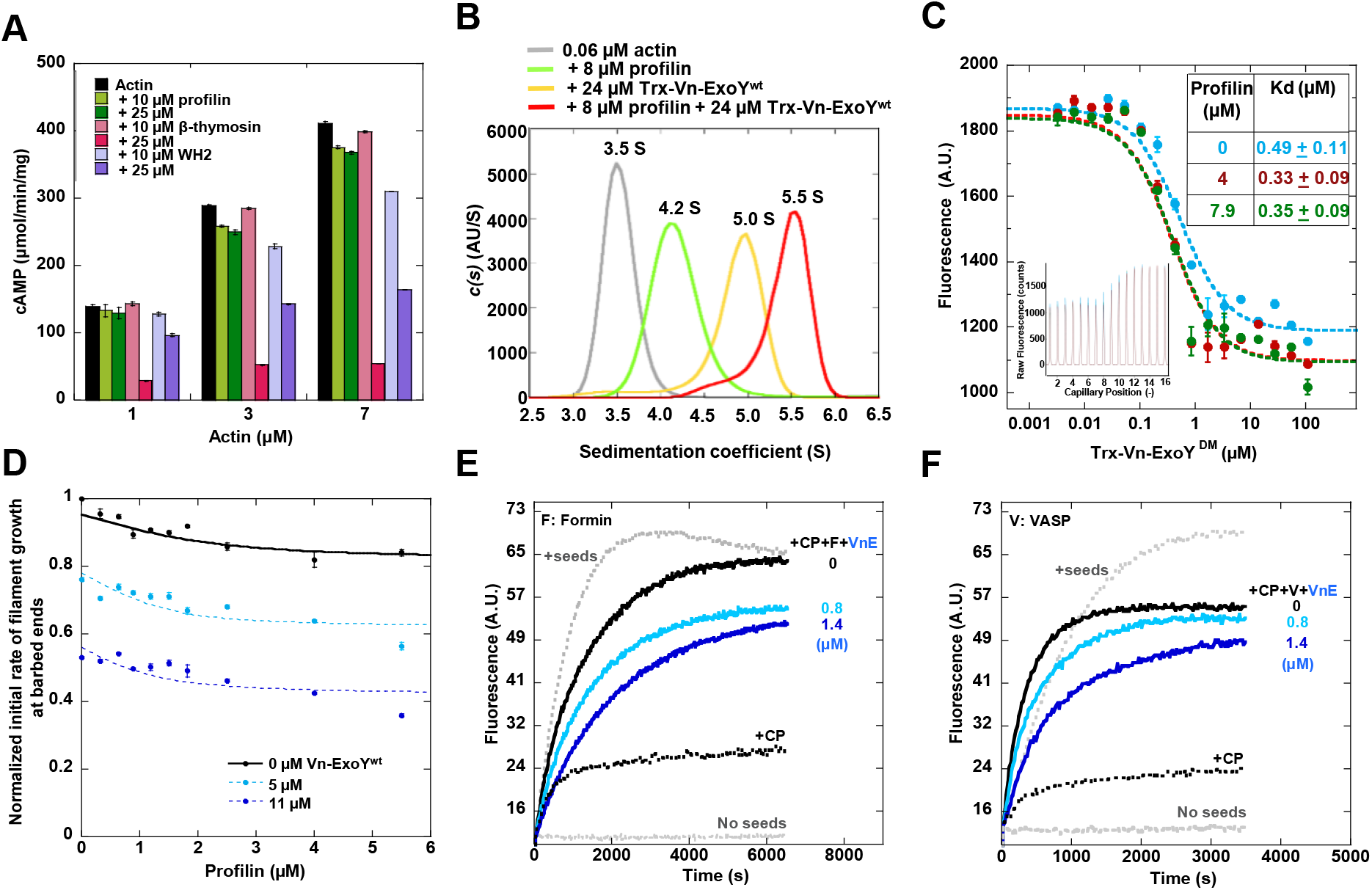
Vn-ExoY is activated by profilin-bound G-actin and inhibits spontaneous or regulated assembly of actin:profilin on F-actin barbed-ends *in vitro*. (A) cAMP synthesis by G-actin-activated Vn-ExoY in the presence of G-ABPs. Reactions containing 5 ng of Vn-ExoY and Latrunculin-A-bound actin at the indicated concentrations (µM) were initiated with 2 mM ATP and incubated at 30°C for 30 min. cAMP synthesis was measured in the absence (black) or presence of two high concentrations (10, 25 µM) of profilin (green), a sequestering β-thymosin (pink) or profilin-like WH2 (purple) domains. The Kd of their interaction with G-actin is 0.1, 0.5 and 0.5 µM, respectively [36, 37]. **(B)** Formation of the Vn-ExoY:actin:profilin complex by AUC-FDS. The c(s) distribution of Alexa-488-labelled-G-actin (60 nM) was determined alone (grey line) and in the presence of either profilin (8 µM; green line), Trx-Vn-ExoY (24 µM; yellow line) or both proteins (red line). **(C)** Measurement of Trx-Vn-ExoY^DM^ binding to free or profilin-bound actin from fluorescence changes in low ionic strength MST experiments. Titration of Alexa-488-labelled, Latrunculin-A-bound ATP-G-actin (0.075 µM) alone (cyan) or bound to 4 (red) and 7.9 (green) µM profilin by Trx-Vn-ExoY^DM^ for a 2-fold dilution series from 108 to 0.033 µM of Trx-Vn-ExoY^DM^ in MST-optimised buffer (see Methods). The inset shows the fluorescence changes in the raw data. The table shows the resulting Kd. Error bars are s.d. (n≥3). **(D)** Vn-ExoY inhibits barbed-end elongation by actin:profilin. Initial barbed-end growth rates from spectrin-actin seeds (0.36 nM) were measured as in figure 1D using 2 µM G-actin (5 % pyrenyl-labelled) and increasing concentrations of profilin in the absence and presence of Vn-ExoY at the indicated concentrations. **(E-F)** Vn-ExoY inhibits formin-(E) or VASP-mediated (F) barbed-end elongation from actin:profilin. Barbed-end growth from spectrin-actin seeds (0.13 nM), 1 µM G-actin (5 % pyrenyl-labelled), and 8 µM profilin was measured in the absence or presence of 2.1 nM of capping protein (CP), 3.2 nM of FH1-FH2 domains of mDia1 formin (denoted by F) (E) or 0.22 µM of VASP (V) (F) at the indicated concentrations of Vn-ExoY (VnE).

### Assembly of actin:profilin at F-actin barbed-ends is inhibited by Vn-ExoY

To understand the effect of Vn-ExoY interaction with actin:profilin on actin self-assembly, we first examined its effect on the spontaneous assembly of actin:profilin at F-actin barbed-ends. We measured the normalised initial rate of barbed-end filament growth in the presence of increasing concentrations of profilin with or without fixed concentrations of Vn-ExoY. Without Vn-ExoY, barbed-end growth was only modestly inhibited by saturating concentrations of profilin as actin:profilin can associate with and elongate F-actin barbed-ends (Fig 2D). With Vn-ExoY, the dependence of barbed-end growth on profilin concentration shifted to lower values in a Vn-ExoY dose-dependent manner. This shift was consistent with a decrease in the G-actin pool available for barbed-end elongation due to its sequestration by Vn-ExoY. We next examined the effect of Vn-ExoY on actin:profilin assembly at F-actin barbed-ends regulated by VASP or formin processive barbed-end elongation factors. The latter bind to barbed-ends via an F-actin binding domain (called Formin-Homology domain 2 (FH2) in formins) and favour actin:profilin recruitment to barbed-ends through their adjacent profilin-binding proline-rich motifs (PRMs, which are repeated in formin FH1 domain). Formins or VASP also counteract the barbed-end capping and growth inhibition by capping proteins (CP). The barbed-end growth of F-actin seeds was therefore monitored in the presence of both G-actin:profilin, CP and either the C-terminal FH1-FH2 domains of mouse Dia1 formin (F or mDia1-FH1-FH2, Figure 2E) or full-length human VASP (V, Figure 2F) to approach the physiological regulation of F-actin barbed-ends. When actin (1µM) is saturated with profilin (8 µM), actin nucleation is prevented, and polymerisation does not occur (horizontal dashed grey lines). Under these conditions, the addition of 0.35 nM of spectrin-actin seeds induced rapid elongation of the barbed ends of actin filaments (ascending dotted grey curves). In the presence of 2.1 nM CP, this barbed-end elongation was partially inhibited (black dashed curves), but restored by the addition of 3.2 nM mDia1-FH1-FH2 (Fig 2E, black curves) or 0.22 µM VASP (Fig 2F, black curves), as both elongation factors compete with CP at F-actin barbed ends. Further addition of Vn-ExoY at 0.8 or 1.4 µM to mDia1-FH1-FH2 or VASP in the presence of CP inhibited the barbed-end growth from actin:profilin induced by the two elongation factors (blue curves). These inhibitory effects were independent of Vn-ExoY AC activity for the duration of the kinetic experiments (S2 Fig). Thus, the interaction of Vn-ExoY with G-actin:profilin inhibits the spontaneous or VASP-/formin-mediated assembly of this complex at the most dynamic ends of F-actin.

### Crystal structures of nucleotide-free and 3’-deoxy-ATP-bound Vn-ExoY in complex with actin-ATP/ADP:profilin or actin-ATP

Next, we investigated the structural mechanism of Vn-ExoY activation by free or profilin-bound G-actin. To improve the crystallisation strategy (see Materials and Methods), we designed a chimeric protein containing three fused functional domains: Vn-ExoY extended at the C-terminus (Vn-ExoY^3455-3896^, either wild-type or with K3528M/K3535I), a short profilin-binding proline-rich motif (PRM) and profilin (Fig 3A, VnE/VnE^DM^-PRM-Prof). The chimera sequestered G-actin with approximately 100-fold higher affinity than the Vn-ExoY^3412-3873^ module and exhibited strong actin-dependent AC activity at low G-actin concentrations (Figs 3B-C). Thus, the actin-binding regions in Vn-ExoY and profilin appear to be fully functional in this chimera. Using this chimera (wild-type or double-mutant), we solved two crystal structures at 1.7 Å resolution: (i) VnE^DM^-PRM-Prof bound to actin-ADP with latrunculin B (LatB), and (ii) VnE-PRM-Prof bound to actin-ADP without LatB (Fig 3D, S1 Table). In these structures, profilin is bound to actin-ADP, the PRM to profilin, Vn-ExoY to actin, while residues 3863-3895 connecting the C-terminus of Vn-ExoY to the PRM are disordered (Fig 3D, S1 Table). Vn-ExoY contacts actin-ADP via its switch A and C, but only switch C is fully ordered. The Vn-ExoY switch A disorder (residues 3682-3727) is close to the LatB-binding site in actin. However, solving the structure without LatB only slightly reduces the switch A disorder. In the latter structure, the NBP of Vn-ExoY is further stabilised by a sulphate ion (Fig 3D, VnE-PRM-Prof-SO_4_^2-^:actin-ADP). On the basis of these structural data, we designed a minimal functional Vn-ExoY module (residues 3455-3863) by removing the C-terminal disordered residues 3864-3895 (Fig 3A). This Vn-ExoY module (shortened by 32 C-terminal residues) shows similar catalytic activity to our original Vn-ExoY^3412-3873^ construct and allowed the determination of two additional crystal structures: (i) nucleotide-free Vn-ExoY bound to actin-ATP-LatB and profilin, with a sulphate ion in Vn-ExoY NBP (Fig 3E, Vn-ExoY-SO_4_^2-^:actin-ATP-LatB:profilin), and (ii) Vn-ExoY bound to actin-ATP-LatB with its NBP filled by a non-cyclisable ATP analogue 3’-deoxy-ATP (3’dATP, Fig 3F), at 1.7 and 2.1 Å resolution, respectively. The Vn-ExoY-SO_4_^2-^:actin-ATP-LatB:profilin structure is very similar to the structures of the VnE/VnE^DM^-PRM-Prof chimera bound to actin-ADP, with the Vn-ExoY switch A similarly disordered. In contrast, the Vn-ExoY-3’dATP-2Mg^2+^:actin-ATP-LatB structure co-crystallised with 3’dATP shows an ordered Vn-ExoY:actin interface. This corresponds to a distinct conformational state of Vn-ExoY NBP that is more competent for catalysis, as described below.

**Fig. 3.**
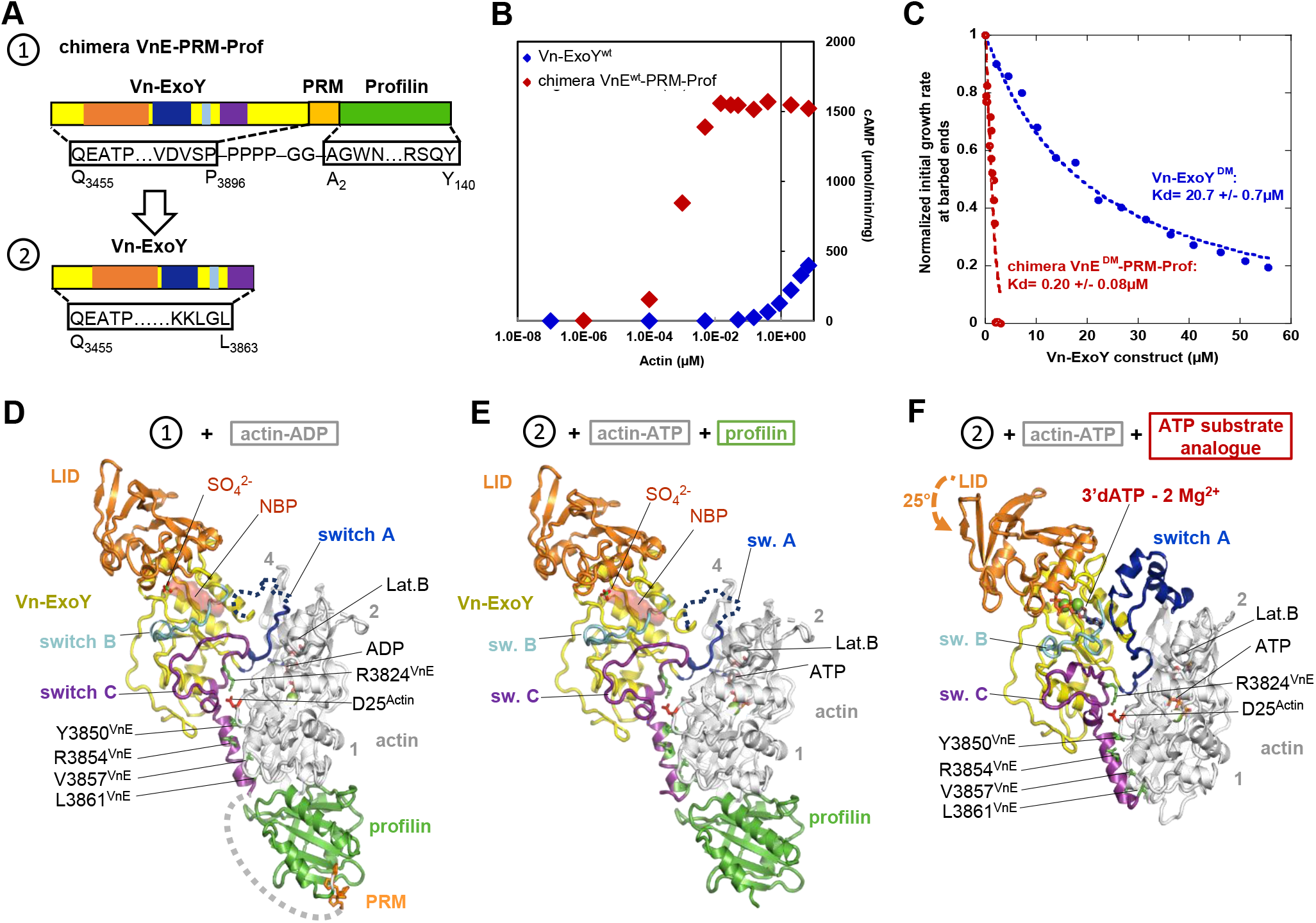
Crystal structures of Vn-ExoY in different conformational states bound to actin:profilin or actin, starting with a chimeric protein containing three fused functional domains including Vn-ExoY. **(A)** Domain diagrams of: (1) the VnE/VnE^DM^-PRM-Prof chimera used to initiate the crystallographic study containing Vn-ExoY extended at the C-terminus (ExoY^3455-3896^, numbering from Uniprot accession no. (AC) F0V1C5) fused to a short PRM/Proline-Rich-Motif capable of binding to profilin, and human profilin-1 (Uniprot AC P07737), and (2) an optimised shorter Vn-ExoY^3455-3863^ construct that remains as active as our original construct Vn-ExoY^3412-3873^ [3]. **(B)** Synthesis of cAMP by actin-activated Vn-ExoY or VnE-PRM-Prof. VnE-PRM-Prof exhibits strong actin-dependent AC activity that saturates at much lower G-actin concentrations. Reactions containing 5 ng Vn-ExoY or 2 ng VnE-PRM-Prof and actin at the indicated concentrations were initiated with 2 mM ATP and incubated at 30°C for 10 or 30 min. **(C)** VnE^DM^-PRM-Prof sequestered G-actin-ATP with 100-fold higher affinity than Vn-ExoY^DM^ (residues 3412-3873) in the barbed-end elongation. Initial barbed-end growth rates from spectrin-actin seeds were measured using 2 µM actin (3 % pyrenyl-labelled) and increasing concentrations of VnE^DM^-PRM-Prof (in red) or Vn-ExoY^DM^ (in blue). **(D)** Structure of the nucleotide-free VnE^DM^-PRM-Prof chimera bound to actin-ADP-LatB. In Vn-ExoY, the C_A_ subdomain and the flexible switch A, switch B, switch C and LID/C_B_ subdomain are shown in yellow, blue, cyan, purple and orange, respectively, here and in all subsequent figures. Disordered regions are shown as dotted lines, the position of the ATP substrate in Vn-ExoY NBP by its surface in red and the four actin subdomains by numbers in grey. Conserved interactions corresponding to hotspots are indicated by black labels, with the Vn-ExoY and actin side chains shown as green and red sticks, respectively. The sulphate ion bound in the VnE-PRM-Prof-SO_4_^2-^:actin-ADP structure is overlaid. **(E)** Structure of nucleotide-free Vn-ExoY in complex with actin-ATP-LatB and profilin (Vn-ExoY-SO_4_^2-^:actin-ATP-LatB:profilin). **(F)** Structure of 3’dATP-bound Vn-ExoY in complex with ATP-actin-LatB (Vn-ExoY-3’dATP-2Mg^2+^:actin-ATP-LatB).

### In the G-/F-actin-bound state, the switch-C of ExoYs ensures stable interactions while other regions remain dynamic

Vn-ExoY undergoes significant structural rearrangements at both the C_A_ and C_B_ subdomains between its nucleotide-free conformation bound to actin-ATP/ADP and profilin, and its 3’dATP-bound conformation in complex with actin-ATP (Fig 4A). At the Vn-ExoY:actin interface, none of the buried contacts of the Vn-ExoY switch C move, except for L3827, which is relocated by the remodelling of switch A (Figs 4A-B). Thus, switch C appears to be essential for initiating the interaction and stabilising Vn-ExoY on actin. In the absence of 3’dATP, most of switch A is too flexible to be modelled, and the switch A portions that are ordered in the absence of ligand undergo profound rearrangements upon 3’dATP-binding (up to 10 Å on D3682 and K3683 Cα atoms, Figure 4A). Subsequently, the letters after the residue numbers are used to distinguish residues of different proteins or functional regions (e.g. ^VnE-CA^ refers to residues belonging to the Vn-ExoY subdomain C_A_). The different conformations of Vn-ExoY bound to actin/actin:profilin with or without a nucleotide substrate analogue allow us to define the mobile “switch” regions as follows: Vn-ExoY switch A as residues Y3675-V3727^VnE-Switch-A^ (I222-N264^PaE-Switch-A^, UniProt AC Q9I1S4_PSEAE), switch B as H3753-A3770^VnE-Switch-B^ (H291-A308^PaE-Switch-B^), switch C as I3819-L3863^VnE-Switch-C^ (K347-V378^PaE-Switch-C^). Other important conformational changes near the interface occur in the parts of switch C in contact with switch A and B, and in the parts of switch B in contact with switch A and C (Figs 4A-B). These regions show high B-factors in all the complexes with nucleotide-free Vn-ExoY (Fig 4D-1). Near the interface, the C_B_ subdomain of Vn-ExoY undergoes the most drastic global rearrangement with a large 25° rotation as a rigid body around the hinge regions 3532-3533^VnE^ and 3662-3663^VnE^. It thus acts as a lid (hereafter referred to as C_B_/LID for residues K3533-M3662^VnE-LID^, corresponding to K86-M209^PaE-LID^ in Pa-ExoY) that closes on C_A_ to stabilise 3’dATP binding (Figs 4A-B, movie S1A-B). The conformational changes induced by 3’dATP binding bring several regions of the Vn-ExoY C_A_ subdomain closer to actin, increasing the buried solvent-accessible area (BSA) of switch A from 338 to 707 Å^2^, reaching the BSA value of switch C (810 and 766 Å^2^, respectively). Vn-ExoY binds to actin over a large surface area, with switch A binding primarily to actin subdomains 2 and 4 and switch C binding to actin subdomains 1 and 3 (Fig 4C).

**Fig. 4.**
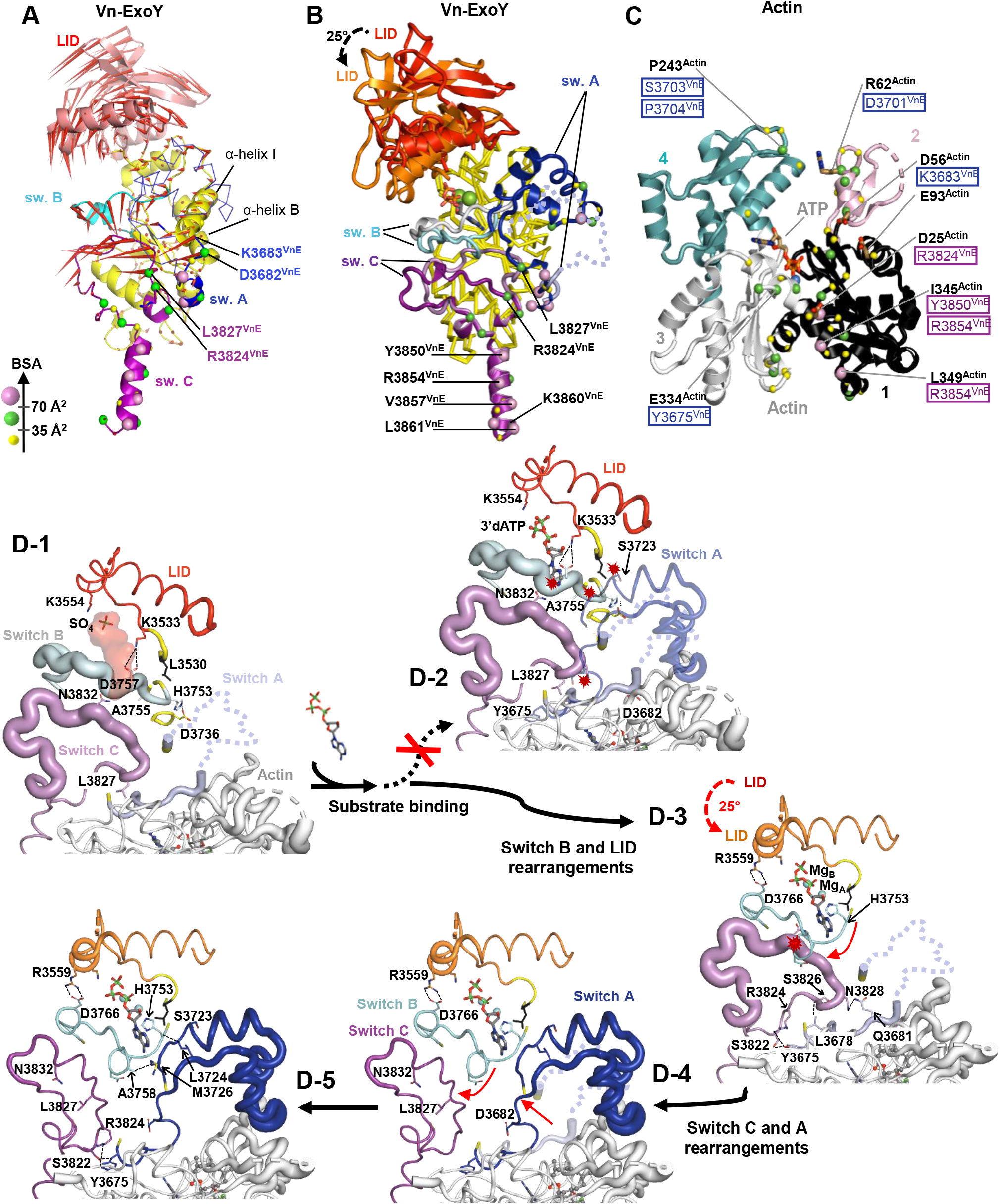
Sequence diagram of the structural rearrangements in actin-bound nucleotide-free Vn-ExoY upon substrate binding. **(A)** Porcupine plot illustrating the direction of Cα atom movement [82] between the nucleotide-free and 3’dATP-bound Vn-ExoY conformations. Red arrow length proportional to Cα atom motion amplitude. **(B)** Vn-ExoY conformations in the Vn-ExoY-SO_4_^2-^:actin-ATP-LatB:profilin and Vn-ExoY-3’dATP-2Mg^2+^:actin-ATP-LatB structures (actin and profilin omitted) overlaid on their C_A_ subdomain. Residues with significant solvent-accessible areas (BSAs) buried by actin are indicated by yellow (BSA ≤ 35 Å^2^), green (between 35 and 70 Å^2^) and pink (≥ 70 Å^2^) spheres on Cα. **(C)** Vn-ExoY-buried actin residues detailing contacts between proteins. The four numbered actin subdomains are shown in different colours. Vn-ExoY residues are boxed with the same colour code as in (A,B). **(D)** The LID/C_B_, switch A, B and C regions of Vn-ExoY are coloured red, light blue, grey and violet in Vn-ExoY-SO_4_^2-^:actin-ATP-LatB:profilin, and orange, blue, cyan and purple in Vn-ExoY-3’dATP-2Mg^2+^:actin-ATP-LatB, respectively. D-1) In the nucleotide-free state, switch B and the beginning of switch C, which contacts switch A and B, are ordered but show relatively high B-factors in Vn-ExoY. The structures are drawn in a cartoon putty representation [82], where the backbone thickness is proportional to the B-factor value. The black dotted lines represent intramolecular interactions. D-2) The conformations of switch B and C, and residues (L3530-F3531^VnE^) near the LID hinge region V3532-K3533^VnE^ are incompatible with 3’dATP binding or the final stabilised conformation of switch A induced by 3’dATP binding. To avoid steric clashes (red explosion symbols), they have to be rearranged. D-3) Structural rearrangement of switch B and LID. LID closure is stabilised by electrostatic bonds between K3535^VnE-LID^ or R3559^VnE-LID^ and D3766^VnE-switch-B^ side chains. The red arrows indicate how the different regions move. D-4) Structural rearrangements of switch A and C. D-5) In this 3’dATP-bound state, switch B and C have B-factors of the same order as those in the overall structure, and the entire switch A stabilises on G-actin. Only the region of switch A (residues 3686-3719) that binds between actin subdomains 2 and 4 retains relatively high B-factors.

To validate the importance of switch C for proper positioning of ExoY on actin, we mutated conserved buried residues of switch C that remain well anchored and stable in the complexes with or without 3’dATP. Actin D25 (D25^Actin^) forms a buried salt bridge at the interface core with the R3824^VnE-switch-C^ side chain (distances of 2.7 and 3.2 Å between N-O atom pairs, Figs 3D-F, 4B-C) and was identified as a critical residue for Pa- and Vn-ExoY activation by F- and G-actin, respectively [3]. The R3824A^VnE-switch-C^ mutation alone results in a 130-fold reduction in Vn-ExoY G-actin stimulated AC activity (Fig 5A). The extreme C-terminus of Pa-ExoY is also critical for its binding to F-actin [38]. The corresponding region in Vn-ExoY is located at the edge of the Vn-ExoY:actin interface, where the extreme C-terminal amphipathic helix of Vn-ExoY (residues 3847-3863^VnE-switch-C^) binds into the hydrophobic cleft between actin subdomains 1 and 3. Mutation of the most buried hydrophobic residues of this amphipathic helix to alanine results in a 240-fold reduction in its AC activity (Fig 5A). Similar interactions occur in the switch C of F-actin-bound Pa-ExoY [27]. Three basic amino-acids of Pa-ExoY near D25^Actin^ may play the same role as R3824^VnE-switch-C^: K347^PaE-switch-C^, H352^PaE-switch-C^ and R364^PaE-switch-C^. While the R364A mutation in Pa-ExoY switch C had almost no effect on its F-actin-stimulated GC activity, the K347A and H352A mutations caused a ∼15- and ∼8-fold decrease, respectively (Fig 5C), indicating that these two amino-acids together function as R3824^VnE-switch-C^. On the other hand, changing the three bulky hydrophobic side chains of the C-terminal amphipathic helix of Pa-ExoY (i.e. F367-L371-F374, Figure 5B) into alanine caused a ∼12,000-fold reduction in F-actin stimulated GC activity (Fig 5C). These residues in the switch C of Vn-ExoY or Pa-ExoY are conserved in most of their counterparts in other β- and γ-proteobacteria (Fig 5D). Thus, when these enzymes switch to their catalytic conformation, their complex with G-/F-actin first stably docks to their cofactor at the switch-C, while other regions remain dynamic to regulate additional functional rearrangements.

**Fig. 5.**
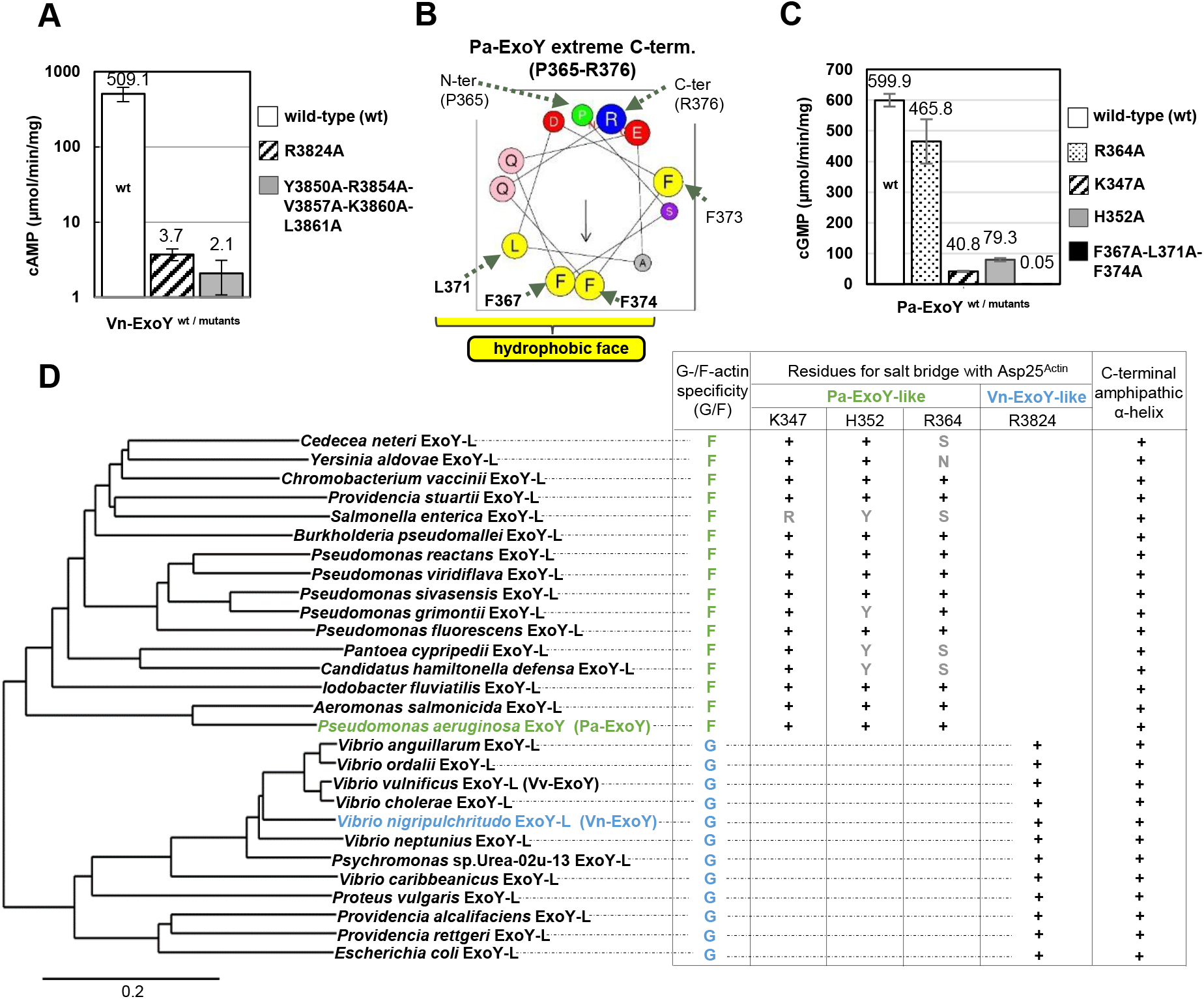
Importance of the switch C at the actin:Vn-ExoY interface. **(A)** cAMP synthesis catalysed by wild-type or switch C mutants of Vn-ExoY (Vn-ExoY^3455-3863^) activated by G-actin. Reactions containing 2.5 or 5 ng for Vn-ExoY and 5, 50 or 100 ng for Vn-ExoY mutants and 3 µM G-actin-ATP-LatB were started with 2 mM ATP and incubated at 30°C for 15 or 30 min. Values are averages of reactions with the indicated enzyme concentrations. **(B)** The amphipathic α-helix at the extreme C-terminus of Pa-ExoY (residues 365-376^PaE-switch-C^) is shown in a helical wheel projection using the HeliQuest analysis (http://heliquest.ipmc.cnrs.fr/). **(C)** cGMP synthesis catalysed by wild-type Pa-ExoY or switch C mutants activated by F-actin. Reactions containing 5 ng Pa-ExoY^wt^ or Pa-ExoY(R364A, K347A, or H352A) or 1 µg Pa-ExoY(F367A-L371A-F374A) and 3 µM F-actin were started with 2 mM GTP and incubated at 30°C for 10 min. Error bars in (A,C) correspond to the s.d. of at least two experimental replicates. In F-actin:Pa-ExoY structure [27], the K347, H352 and R364 side chains are at distances of 3.1, 4.5 and 4.1 Å from the D25^Actin^ side chain, respectively. **(D)** Phylogenetic tree showing the relationships between ExoY-like proteins/effector domains produced by different β- and γ-proteobacteria. Vn- and Pa-ExoY are representative of the distant ExoYs. ExoYs are classified as Vn- or Pa-ExoY-like homologues based on both their sequence similarity and the predicted complex with actin by AlphaFold [84] (see Fig. S10 for more details). A + sign indicates that the Vn- or Pa-ExoY residue/structural element is conserved at the same position in the ExoY-like homologue’s sequence and predicted structure.

### Significance of switch A and C_B_/LID closure in stabilising nucleotide and Mg^2+^

Apart from a difference in the structural topology of the two ExoY proteins (S3 Fig), the overall structure of Vn-ExoY bound to actin and 3’dATP is close to that of *V. vulnificus* Vv-ExoY bound to actin:profilin [27] (S4A Fig). In the 3.9-Å resolution cryo-EM structure, the 3’dATP ligand used to prepare the complex could not be modelled in Vv-ExoY. In both structures, switch A adopts a similar conformation between actin subdomains 2 and 4. This interaction primarily prevents Vn- and Vv-ExoY from binding along the side of F-actin. This explains their G-actin specificity, whereas the orientation of Pa-ExoY switch A bound to F-actin differs [27] (S4B Fig). Furthermore, the positioning of the Vn-ExoY switch A on G-actin prevents the association of Vn-ExoY:G-actin or Vn-ExoY:G-actin:profilin complexes with F-actin barbed-ends (S5A Fig) and is responsible for the inhibition seen in Figure 1D, 2D-F. Switch A is also the region with the most divergent folding at the toxin:cofactor interface between G/F-actin-bound ExoYs and CaM-bound EF or CyaA (S5B-C Figs). Thus, this region appears to be evolutionarily essential for the cofactor specificity of NC toxins.

Our different structures reveal two major consecutive conformational changes occurring at the catalytic site of NC toxins. In the free/inactive Pa-ExoY structure [25], the A, B, and C switches were not visible as partially degraded and/or flexible. In our nucleotide-free structures of Vn-ExoY bound to G-actin-ATP/-ADP and profilin (Figs 3D-E), the Vn-ExoY NBP is still only partially stabilised by cofactor binding (Fig 4D-1), showing that cofactor binding is not sufficient to fully stabilise the NBP and catalytic residues of NC toxins, as previously suggested [2, 23, 27]. 42 to 46 of the 53 residues from switch-A are still disordered. A SO_4_^2-^ ion is bound to the Vn-ExoY NBP via residues (K3528^VnE-CA^, K3535^VnE-LID^, S3536^VnE-LID^, K3554^VnE-LID^) that coordinate the γ- and β-phosphates of 3’dATP when this ligand is bound (Fig 4D-2). Binding of the ATP substrate and Mg^2+^ ion in the Vn-ExoY NBP requires rearrangement of switch B and the supporting part of switch C (Fig 4D-3). To reach its most favourable binding position, the nucleotide must move ∼4 Å (distance between the SO_4_^2-^ ion and 3’dATP γ-phosphate) through the LID closure (Fig 4D-3). Structural rearrangements in the region of switch C, which supports switch B, are accompanied by stabilisation of switch A and the substrate analogue (Figs 4D-4,4D-5). In the final stage, switch B and C are well-ordered, while the entire switch A is stabilised on G-actin (Fig 4D-5). In the Vn-ExoY-3’dATP-2Mg^2+^:actin-ATP-LatB structure, switch A ultimately contributes to the stabilisation of the nucleotide base through interactions with M3726^VnE-switch-A^

### Catalytic mechanism of ExoY NC toxins activated by G- or F-actin with purine nucleotides

In the known cofactor:toxin structures, the nucleotide ligands adopt different positions and conformations in the toxin catalytic pocket, variably coordinated by one or two metal ions (S6-S7 Figs). The Pa-ExoY-bound 3’dGTP is coordinated by a single metal ion in the F-actin-activated Pa-ExoY structure [27]. In the structure of Vn-ExoY bound to actin and 3’dATP, the electronic density and coordination geometry suggest that 3’dATP is coordinated by two Mg^2+^ ions. To unambiguously distinguish metal ions from water, we replaced Mg^2+^ with Mn^2+^ ions and collected an anomalous data set at the manganese absorption edge (see Materials and Methods). The anomalous difference Fourier map confirms that two Mn^2+^ ions are bound to 3’dATP at the positions occupied by the Mg^2+^ ions, with no significant changes elsewhere (Fig 6B). Mg^2+^_A_ is pentacoordinated in a square pyramidal geometry by two aspartates D3665^VnE-CA^ and D3667^VnE-CA^, and H3753^VnE-switch-B^, which are invariant in all NC toxins (Fig 6A), an α-phosphate oxygen of 3’dATP, and a water molecule. Mg^2+^_B_ is octahedrally hexacoordinated by the same two aspartates (D3665/D3667^VnE-CA^), one oxygen from each of the three phosphates of 3’dATP and a water molecule.

**Fig. 6.**
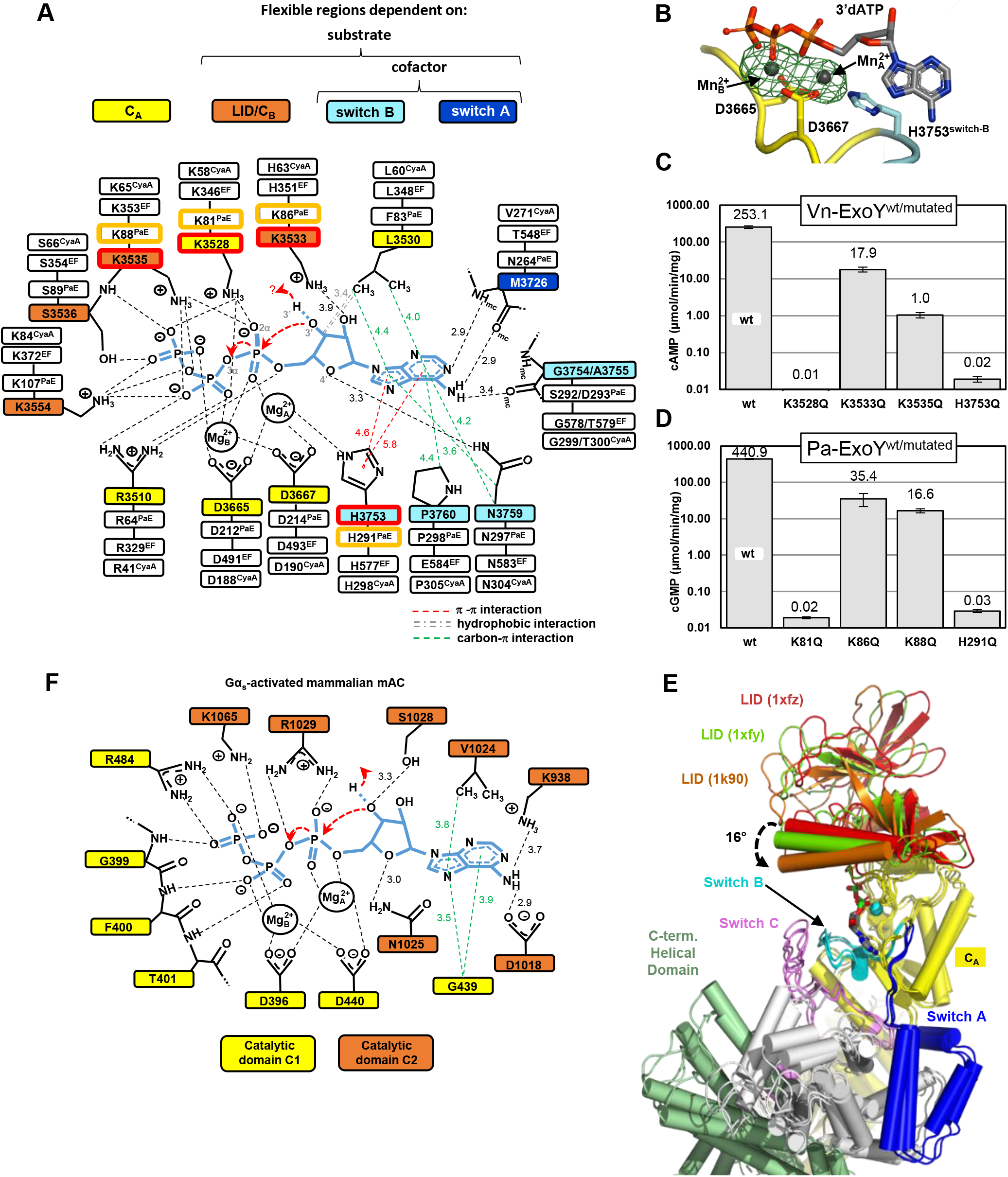
Model of ATP-binding in activated Vn-ExoY and mammalian transmembrane AC (tmAC) and residues important for the purinylyl cyclase activities of Vn- and Pa-ExoY. **(A)** Model of ATP-binding in activated-Vn-ExoY based on its interactions with 3’dATP in the Vn-ExoY-3’dATP-2*Mg^2+^:ATP-actin-LatB structure. Thresholds used for interaction detection are those of the PLIP [85] and Arpeggio [86] web servers and the PoseView tool in the ProteinsPlus web server [87]. **(B)** The model-phased anomalous difference Fourier map from data collected at 6.55 keV, shown as a green mesh contoured at 5.0 σ, confirms the presence of two Mn^2+^ (grey spheres) bound to 3’dATP in Vn-ExoY NBP. **(C)** cAMP synthesis catalysed by wild-type or mutants of Vn-ExoY (Vn-ExoY^3455-3863^) activated by G-actin. Reactions containing 10 ng Vn-ExoY, 100 ng Vn-ExoY(K3533Q) or 1 µg Vn-ExoY(K3528Q, K3535Q, or H3753Q) and 3 µM of LatB-Mg-G-actin were started with 2 mM ATP and incubated at 30°C for 10-30 min. **(D)** cGMP synthesis catalysed by wild-type or mutants of Pa-ExoY activated by F-actin. Reactions containing 10 ng Pa-ExoY^wt^/Pa-ExoY(K86Q or K88Q) or 8 µg Pa-ExoY(K81Q or H291Q) and 3 µM of Mg-F-actin were started with 2 mM GTP and incubated at 30°C for 10-30 min. **(E)** The LID/C_B_ subdomain of EF occupies different positions relative to the C_A_ subdomain between the nucleotide-free (1xfz, 1xfy) [26] and nucleotide-bound (1k90) [23] structures of CaM-bound EF. It rotates between the 1xfz and 1k90 PDBs by 16° as a rigid body. The structures are superimposed on their C_A_ subdomains shown in yellow. **(F)** Model of ATP-binding in activated class III tmAC based on the structure of the type V AC C1a/type II AC C2 heterodimer bound to the ATP analogue RP-ATPαS (PDB 1CJK) [18]. Activated Vn-ExoY and tmAC closed NBPs have similar interactions with ATP analogues. For example, their conserved asparagine N3759^VnE-switch-B^ or N1025^tmAC-IIC2^ similarly hydrogen-bonds the ribose ring O4’-oxygen of ATP analogues. In tmAC, the interaction of N1025^tmAC-IIC2^ with ATP is essential for correct ribose positioning during catalysis [18, 42].

Figure 6A summarises the main interactions observed with 3’dATP in the Vn-ExoY-3’dATP-2*Mg^2+^:actin-ATP-LatB structure, extrapolated to the natural substrate ATP. The residues of Vn-ExoY that coordinate the nucleotide and metal-ions are indicated in coloured boxes in the first inner shell. The corresponding residues in the Pa-ExoY, EF, and CyaA sequences are in the next outer shells. 9 out of the 14 substrate and Mg^2+^ ion binding residues belong to regions of the enzyme that are flexible and significantly rearranged and stabilised upon binding of the cofactor, nucleotide substrate and metals. This explains the low basal activity of the free enzyme [5]. All side-chain contacts are provided by residues that are invariant or similar in Pa-ExoY, EF, and CyaA, and most interactions are conserved in CaM-activated EF bound to 3’dATP with a Yb^3+^ ion and a closed LID (PDB 1k90) [23] (S7D Fig).

To assess the relevance of the observed interactions with the ATP analogue, some of the conserved residues involved were mutated in Vn- and Pa-ExoY and the effect on their AC and GC activity was investigated. H351^EF-LID^ in EF has been proposed to play a role in the 3’OH deprotonation, either by accepting the 3’OH proton (general base), or by facilitating the presence of OH^-^ ions near the ATP 3’OH group [23, 26, 29, 31, 32, 39]. H351^EF-LID^ is conserved in CyaA, but in ExoYs it corresponds to a lysine: K3533^VnE-LID^ or K86^PaE-LID^ (Fig 6A). In the active site of Vn-ExoY bound to actin, 3’dATP and 2 Mg^2+^ or 2 Mn^2+^, the K3533^VnE-LID^ side chain is too distant to potentially coordinate the ATP 3’OH moiety if the latter were present. K3533Q^VnE-LID^ or K86Q^PaE-LID^ mutations only decreased G/F-actin-activated AC and GC activity by 14- and 12-fold, respectively (Figs 6C-D), further suggesting that these residues are unlikely to act as a general base. Next, we examined K3528^VnE-CA^ and K3535^VnE-LID^, which are close to the bridging O3α and non-bridging O2α, O1β and O2γ oxygens of 3’dATP (Fig 6A). Both side chains are ∼3.4 Å from O3α. K3528^VnE-CA^ is closer to O2α and O2γ (2.8-2.9 Å) than K3535^VnE-LID^, and K3535^VnE-LID^ is closer to O1β (3.3 Å). The catalytic activity of Vn- and Pa-ExoY was essentially abolished (at least ∼23,000-fold reduction) by the K3528Q^VnE-CA^ and the corresponding K81Q^PaE-CA^ mutations in Pa-ExoY. The K3535Q^VnE-LID^ or the equivalent K88Q^PaE-LID^ mutations resulted in a 250- and 26-fold decrease in G-/F-actin-induced Vn-ExoY AC or Pa-ExoY GC activity, respectively (Figs 6C-D). The invariant H3753^VnE-switch-B^, which coordinates Mg^2+^_A_ and stabilises the purine base via a π-π stacking with its imidazole ring, is specific to NC toxins. Mg^2+^_A_ may be particularly important in the NBP of activated mammalian ACs to coordinate and polarise the 3’OH of ATP, facilitating its deprotonation as a first step in the cyclisation reaction [18, 19, 40]. The H3753Q^VnE-switch-B^ or equivalent H291Q^PaE-switch-B^ mutations greatly reduced the catalytic activity of the toxins (∼13,000-fold reduction, Figs 6C-D), suggesting a critical role for the switch-B invariant histidine and Mg^2+^_A_ in the catalysis of NC toxins.

### Structural basis for the pyrimidinyl cyclase activity of ExoYs

The pyrimidinyl cyclase activity of NC toxins has been identified *in vitro* [5, 41] or in eukaryotic cells [4, 15]. On the other hand, no structural characterisation has yet been performed. Among the pyrimidine nucleotides, Vn-ExoY, EF or CyaA all primarily use CTP as a substrate, whereas Pa-ExoY prefers UTP but can also use CTP. To determine how activated NC toxins can use small pyrimidine nucleotides as substrates, we solved the structure of Vn-ExoY bound to 3’dCTP in complex with actin-ADP at 2.2 Å resolution (Vn-ExoY-3’dCTP-2*Mn^2+^:actin-ADP-LatB). To our knowledge, this is the first structure of an activated NC enzyme to be determined in association with a pyrimidine nucleotide substrate analogue. The overall conformation of Vn-ExoY in this structure is identical to those found in the complexes with actin-ATP, 3’dATP and two Mg^2+^ or Mn^2+^. As with 3’dATP, the LID of Vn-ExoY is closed on 3’dCTP and two metal ions coordinate the phosphate moiety of 3’dCTP. In the NBP of Vn-ExoY, 3’dCTP is stabilised by the same residues and types of interactions as 3’dATP (Fig 7A). The main differences are in the interactions with the nucleotide ribose and base moieties. These are stabilised by fewer and weaker hydrogen bonds and framed by fewer hydrophobic interactions with 3’dCTP than with 3’dATP. With 3’dATP, the ribose is stabilised by L3530^VnE-CA^ hydrophobic interactions and hydrogen-bonded by K3533^VnE-LID^ and N3759^VnE-switch-B^ side-chains (Fig 6A). In the activated structures of Vn-ExoY and tmAC [18] with an NBP closed on an ATP analogue, N3759^VnE-switch-B^ and N1025^tmAC-IIC2^ similarly hydrogen-bond the O4’ oxygen of the nucleotide ribose ring (Figs 6A, 6F). Just as mutations at N1025^tmAC-IIC2^ reduce tmAC kcat by 30-100 [42], mutations at N583^EF-switch-B^ in EF (equivalent to N3759^VnE-switch-B^) reduce EF catalytic activity by at least two orders of magnitude [23]. These data highlight the importance of hydrogen bonding with the nucleotide ribose for correct positioning of the substrate during class II and III AC catalysis. With 3’dCTP, electrostatic interactions with the nucleotide ribose switch to a single, more labile hydrogen bond between the cytidine ribose ring O4’-oxygen and a water molecule coordinated by D3756^VnE-switch-B^ or N3759^VnE-switch-B^. The cytosine base is hydrogen-bonded by M3726^VnE-switch-A^ and A3755^VnE-switch-B^ as for the adenine base. However, these residues only coordinate the N4 atom of cytosine. For 3’dATP, they stabilise both the N1 and N6 atoms of adenine through shorter hydrogen bonds. Similarly, in the Pa-ExoY:3’dGTP-1*Mg^2+^:F-actin structure, the N1, N2, and O6 guanine atoms are hydrogen-bonded by the Ser292^PaE-switch-B^, H291^PaE-switch-B^, and Glu258^PaE-switch-A^ side chains, respectively (S7C Fig). The pyrimidine base is otherwise surrounded by similar hydrophobic interactions to the purine base of 3’dATP. However, it lacks Pro3760 side-chain interaction with imidazole ring of 3’dATP purine base on solvent accessible side. On the triphosphate moiety of 3’dCTP, the phosphates and two metal ions are tightly coordinated and less solvent-accessible than the nucleoside moiety, which is very similar to 3’dATP.

**Fig. 7.**
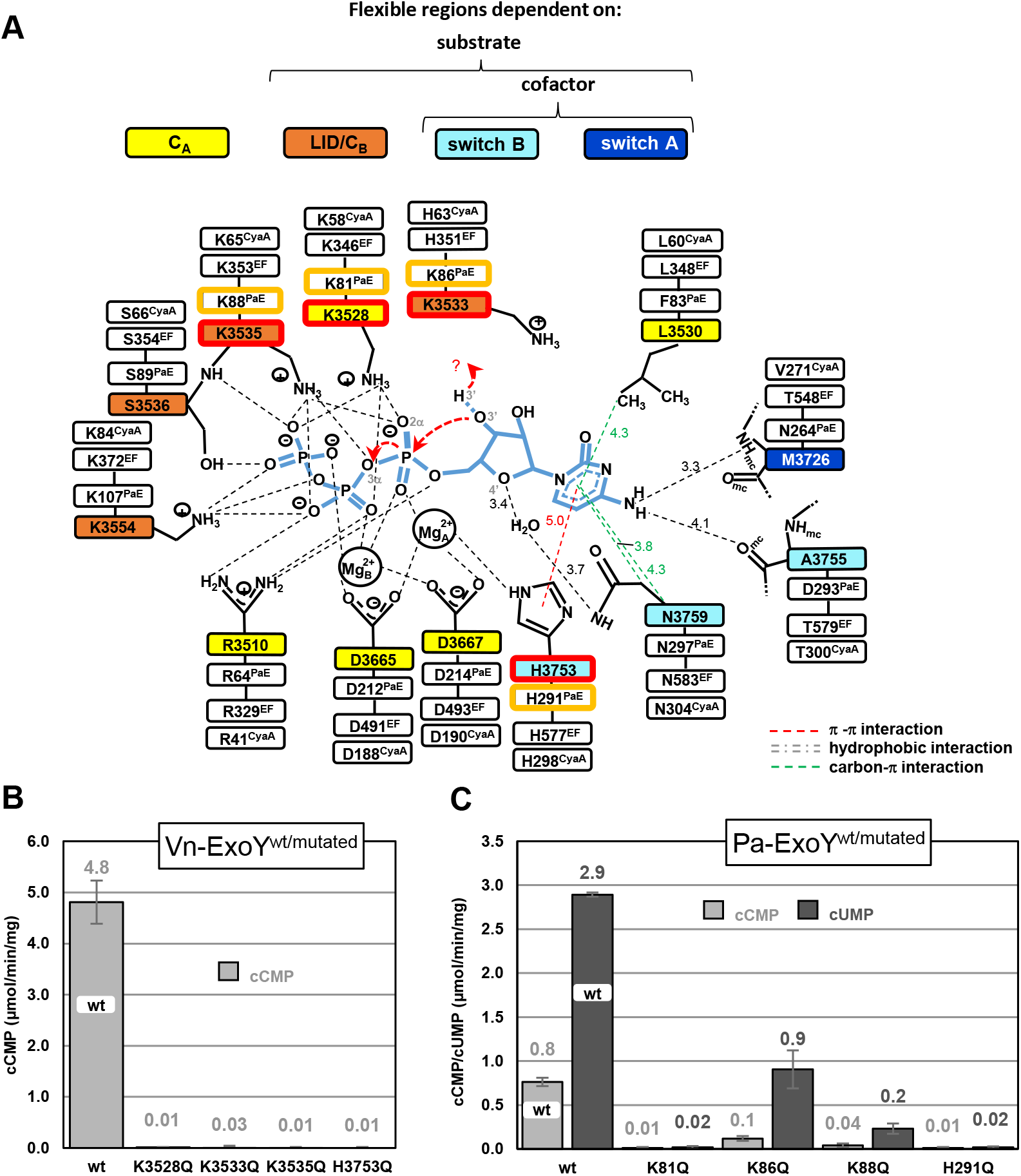
Model of CTP-binding in actin-activated Vn-ExoY and residues important for the CC or UC activity of Vn- and Pa-ExoY. **(A)** Model of CTP-binding in activated-Vn-ExoY based on its interactions with 3’dCTP in the Vn-ExoY-3’dCTP-2*Mn^2+^:ADP-actin structure. Interaction detection thresholds are as in Figs 6A, F. **(B)** cCMP synthesis catalysed by wild-type or mutants of Vn-ExoY activated by G-actin. Reactions containing 100 ng Vn-ExoYwt or 1 µg-100 ng (mean) Vn-ExoY(K3528Q, K3533Q, K3535Q, or H3753Q) and 6 µM of Lat.B-Mg-G-actin were started with 2 mM CTP and incubated at 30° C for 60 min. **(C)** cCMP and cUMP synthesis catalysed by wild-type or mutants of Pa-ExoY activated by F-actin. cCMP synthesis was performed by using 1µg-100 ng (mean) Pa-ExoY^wt^ or Pa-ExoY(K81Q, K86Q, K88Q, or H291Q) and 6 µM of Mg-F-actin and reactions were started with 2 mM CTP and incubated at 30° C for 60 min. cUMP synthesis was performed by using 100-200 ng (mean) Pa-ExoY^wt^ or 1000-100 ng (mean) Pa-ExoY(K81Q, K86Q, K88Q, or H291Q) and 6 µM of Mg-F-actin and reactions were started with 2 mM UTP and incubated at 30° C for 60 min. In the absence of G-/F-actin, the basal CC and UC activity of Vn- and Pa-ExoY are 0.007 and 0.003 pmol/min/ng of the enzyme, respectively.

The mutations analysed above for Vn-ExoY AC or Pa-ExoY GC activity were reconsidered and their effect on their cytidylyl or uridylyl cyclase (CC/UC) activity was examined to determine whether the conserved interactions with the nucleotide triphosphate moiety play a similar role in the purinyl and pyrimidinyl cyclase activity of ExoYs. The mutations with the greatest effect on Vn-ExoY CC or Pa-ExoY UC activity (150- to 480-fold reduction) are those that also most affect their AC or GC activity, namely K3528Q^VnE-CA^ or K81Q^PaE-CA^ and H3753Q^VnE-switch-B^ or H291Q^PaE-switch-B^, respectively (Figs 7B-C). As with 3’dATP, the K3528^VnE-CA^ side chain hydrogen-bonds the α, β, and γ phosphate oxygens of 3’dCTP, and the H3753^VnE-switch-B^ side chain stabilises both Mg^2+^ via a hydrogen bond and the cytosine base via a π-π stacking with its imidazole ring. Taken together, these results demonstrate that NC toxins use a similar two-metal-ion catalytic mechanism with purine and pyrimidine nucleotide substrates, relying on the close coordination of their triphosphate moiety for catalysis.

## Discussion

We have used the MARTX Vn-ExoY module to explore the functional specificities of ExoY-type toxins that selectively interact with G-actin *in vitro* [5]. In cells, G-actin is not free, but bound to cytoskeletal G-actin-binding proteins that prevent uncontrolled G-actin self-assembly. Compared to the Kd of ∼0.2-20 nM for CyaA or EF binding to CaM [33] or of ∼1 µM for Pa-ExoY binding to F-actin [3], Vn-ExoY has only a modest affinity for G-actin (Figs 1D-E, Kd∼12 µM). This is balanced by its ability to interact with and be efficiently activated by G-actin bound to profilin (Figs 2A-C), as the G-actin:profilin complex is abundant in eukaryotic cells [35, 43]. Due to overlapping binding interfaces at the hydrophobic cleft between actin subdomains 1 and 3, Vn-ExoY cannot interact with G-actin in complex with other common G-actin-binding proteins such as Thymosin-β4, WH2 domains (Fig 2A) or actin-depolymerising factor (ADF)/cofilin (S9 Fig). The G-actin:profilin complex serves as a physiological cofactor for Vn-ExoY and its closely-related ExoYs in eukaryotic cells. Our structural bioinformatics analysis suggests that many ExoY-like homologues from different γ-proteobacteria are structurally-related to Vn-ExoY (Fig 5D). The G-actin:profilin complex is important for supporting filament assembly at barbed-ends. Vn-ExoY interaction with G-actin:profilin inhibits both spontaneous and formin- or VASP-mediated association of G-actin:profilin complexes with F-actin barbed-ends *in vitro*, independent of the toxin’s NC activity (Figs 2D-F, S2). As shown by the multipartite interactions of our actin-bound VnE-PRM-Prof chimera, the ternary complex does not prevent profilin from interacting with PRM segments (Fig 3D).

Pathogens using NC toxins have therefore adopted two alternative strategies during evolution by turning to actin-activated NC toxins. Unlike calmodulin, actin self-assembles in a very dynamic manner. To trigger their toxic NC activity, on the one hand, Pa-ExoY-type ExoYs bind along filaments (Belyy et al, 2016), which are particularly abundant beneath the plasma membrane of eukaryotic cells (Koestler et al, 2009). On the other hand, Vn-ExoY/Vv-ExoY-type ExoYs exclusively target monomers in one of their most abundant cellular forms, namely the polymerisation-ready actin pool, the actin:profilin complex. As the turnover of this pool is high at sites of active cytoskeletal remodelling, the interaction interface of these actin-utilising NC toxins has evolved to form a ternary Vn-ExoY:actin:profilin complex that sequesters actin:profilin, prolonging the lifetime of Vn-ExoY or related ExoYs bound to actin:profilin and thus their activation and cytotoxicity. The interaction of Vn-ExoY with actin:profilin may be particularly important in determining subcellular localisation and activation at sites of active actin cytoskeleton remodelling (Fig 8). At these sites, G-actin:profilin and elongation-promoting factors play a prominent role together in fine-tuning the coordination and competition of F-actin networks [44, 45]. This fine-tuning of actin dynamics is coordinated by a large number of multidomain ABPs and is tightly associated with cAMP signalling in important cellular processes, including cell adhesion and T-cell migration [1, 46–48]. VASP activity is regulated by PKA and PKG [49] and is important for immune function [50]. Recruitment of the abortive ternary complex Vn-ExoY:actin:profilin by the PRMs of multidomain ABPs may be of interest to pathogens to confine Vn-ExoY-like toxin activity to these discrete sites where signalling and cytoskeletal remodelling are intense and highly intertwined. These results pave the way for further investigation of the subcellular compartments in which G-actin:profilin-selective ExoYs produce supraphysiological amounts of purine and pyrimidine cyclic nucleotides, and for elucidation of their specific virulence mechanisms compared to F-actin-targeting ExoYs.

**Fig. 8.**
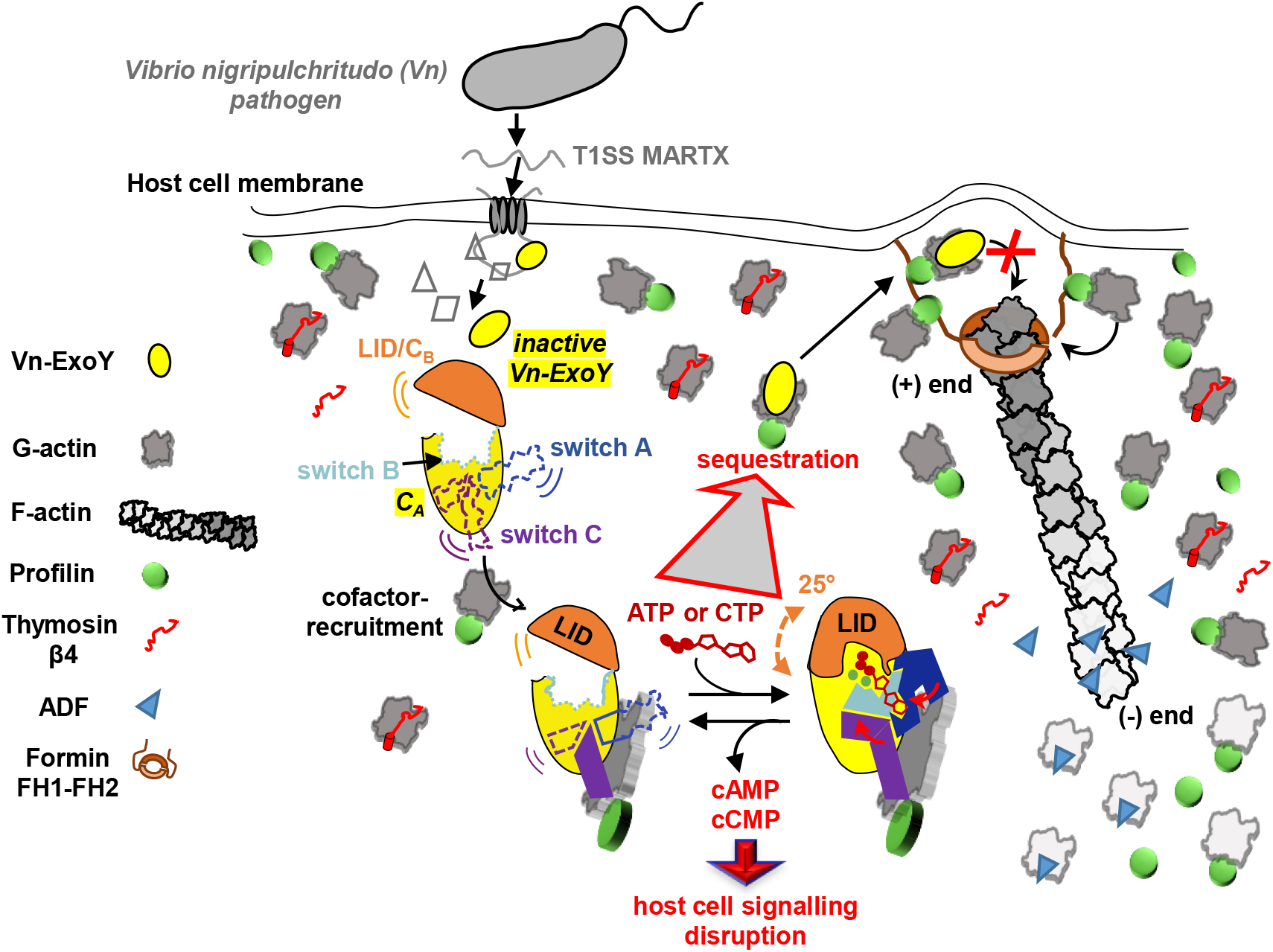
Model of the activation mechanism and cytotoxic effects of Vn-ExoY elicited by its interaction with actin:profilin. **(A)** Once secreted by bacterial T1SS, the MARTX toxin binds to the target cell membrane, translocates and releases its various MARTX effector modules into host cells [14], including Vn-ExoY. **(B)** In isolation, the three switch regions and the CB/LID domain of Vn-ExoY are flexible and unstable, and the enzyme is inactive. **(C)** The interaction of Vn-ExoY with the actin:profilin complex is first tightly docked to actin by switch C. This first step brings together and stabilises only part of the substrate and metal ion binding sites: it initiates stabilisation of switch B and brings switch A closer to the NBP. This intermediate activated conformation allows the active site to ‘breathe’, particularly at the C_B_/LID subdomain of NC toxins, which is close to the cofactor binding site. The C_B_/LID subdomain of NC toxins can therefore continue to adopt a variety of positions. **(D)** Substrate and metal ion binding promotes C_B_/LID closure of NC toxins. This allows the cofactor pre-stabilised active site of NC toxins to accommodate and ultimately constrain all substrate and metal ion interactions. By simply closing and opening the C_B_/LID to load, process nucleotide substrates of different sizes (purine/pyrimidine nucleotides) and release the products, the large substrate-induced fit movement of the NC toxin C_B_/LID subdomain ensures both good substrate specificity and efficient enzymatic turnover. **(E)** Sequestration of G-actin:profilin by Vn-ExoY (or Vn-ExoY-like ExoYs) in an abortive ternary complex, which can be recruited by ABPs through their PRMs (present in formin FH1) but cannot deliver the bound G-actin to the F-actin (+) ends, may help to prolong the lifetime of the Vn-ExoY active state and confine its toxic NC activity to sites of active actin remodelling (shown here as a membrane protrusion initiated by formin elongation factor). At these sites, G-actin:profilin complexes and VASP, Arp2/3 or formin elongation-promoting factors fine-tune the homeostasis of F-actin networks in close cooperation with cAMP-dependent signalling [45–48].

Our structural data show that Vn-ExoY switch C interactions at the Vn-ExoY:actin interface remain stable with different conformations of the LID/C_B_ subdomain and switch A (Figs 3D-F, 4A-B), thus firmly docking the toxin to actin. Our mutational analysis reveals two hotspots in the switch C of Vn- and Pa-ExoY, whose mutation almost completely abolishes their G- or F-actin-dependent NC activity (Fig 5). These sites could be hits in the search for ways to inhibit ExoY toxins. In contrast, switch A is only barely stabilised by actin in the absence of substrate (Figs 3D-E, 4D). EF and CyaA, on the other hand, have a switch A that is already fully folded on CaM in their nucleotide-free, cofactor-bound state [23, 24]. However, their C_B_/LID subdomain can also adopt different positions in isolated CyaA [51] or between different crystal structures of CaM-bound EF (Figs 6E, S8). Many crystal structures of CaM-activated EF or CyaA with nucleotide ligands in the active site were first crystallised without substrate analogues or reaction products and then incubated with these ligands. Crystal packing in these structures may have prevented complete rearrangement of the NBP and LID/C_B_ subdomain closure. As a result, and due to the intrinsic flexibility of the LID/C_B_ subdomain, both substrate analogues and metal ions occupy quite different positions in the NBP of CaM-bound EF or CyaA (S6-S8 Figs) [1]. These structures may correspond to intermediate stages of LID/C_B_ movement and substrate entry. Cofactor binding thus initiates allosteric stabilisation of switch B in NC toxins by repositioning the central switch A and the C-terminal switch C towards switch B, as previously reported [2, 23, 27]. In the ExoY NC toxins, however, complete stabilisation of switch A on actin also requires substrate binding and C_B_/LID closure. Switch A is involved in the correct positioning of the substrate by directly coordinating the purine or pyrimidine base of the nucleotides in all cofactor-bound NC toxins (Figs 6A, 7A, S7). In both the inactive conformation of free Pa-ExoY [25] and the nucleotide-free conformations of actin:profilin-bound Vn-ExoY (Figs 3D-E), H291^PaE-switch-B^ and H3753^VnE-switch-B^, respectively, are stabilised away from the NBP by a salt bridge with D3736^VnE^/D274^PaE^, which are located near the switch A (Fig 4D). After cofactor binding, substrate binding in ExoYs induces the flipping of H3753^VnE-switch-B^/H291^PaE-switch-B^ towards the NBP. This invariant histidine of NC toxin switch B assists in the correct positioning of the catalytic Mg^2+^_A_ on the two invariant aspartates (D3665D3667^VnE-CA^ or D212D214^PaE-CA^) with purine or pyrimidine nucleotide substrate analogues (Figs 6A-B, 7A). Thus, for all NC toxins, substrate binding ultimately facilitates C_B_/LID closure. This increases the enzyme interactions with substrate and metal ions on the one hand, and the interactions between the substrate and the metal ions on the other hand (S7A-E Figs).

Our data demonstrate unambiguously that Vn-ExoY uses a two-metal-ion-assisted catalytic mechanism with purine and pyrimidine nucleotides. Given the sequence similarity and structural conservation, this is most likely to be the case for all bacterial NC toxins after LID/C_B_ closure (Figs 6A-B, 7A, S7). Interestingly, the coordination of metal ions and residues of the Vn-ExoY NBP with the ATP-analogue is also closer to that observed in the structures of activated class III ACs [18] than to that previously observed in the NBPs of cofactor-activated NC toxins (Figs 6A, 6F, S7). The active site of transmembrane or soluble class III ACs (tmAC/sAC) is located at the interface of their two homologous catalytic domains (termed C1 and C2 in tmAC, Figure 6F). It has also been proposed to undergo an open-closed transition during catalysis. In the case of tmAC/sAC, however, the exact role of the open-closed transition of their dimeric NBP and the identification of the dynamic active site processes during substrate binding and turnover remain to be clarified [19]. In the closed NBP of all ACs, the correct positioning of the different ATP moieties is essential for catalysis. This is shown by the strong effect of mutations targeting N1025^tmAC-IIC2^ in tmAC [18, 42] or N583^EF-switch-B^ in EF [23], which corresponds to N3759^VnE-switch-B^ in Vn-ExoY (Figs 6A, 6F). On the phosphate and 3’OH side, the most probable reaction sequence as well as the key interactions during each step were investigated by computational studies for tmAC with a closed ATP-binding NBP [40] and for EF with an open or nearly-closed ATP-binding NBP (using PDB 1XFV/1SK6) [31]. The reaction model for tmAC better fits the numerous and tight interactions around 3’dATP and metal ions observed in the closed NBP of activated Vn-ExoY. The proposed mechanism for tmAC is a substrate-assisted general base catalysis, where several residues of the enzyme appear to be particularly important with metal-ion cofactors for lowering the two highest energy barriers of the reaction. First, K1065^tmAC-IIC2^, Mg^2+^_A_, Mg^2+^_B_ and the two conserved D396D440^tmAC-VC1^ anchoring metal ions are critical for jointly facilitating the proton transfer from the ribosyl 3′O to a γ-phosphate oxygen [40]. R1029^tmAC-IIC2^ is then critical for the concerted phosphoryl transfer step in which K1065^tmAC-IIC2^ is also involved. These interactions belong to C1 and C2 structural elements that are distant in the open NBP of tmAC [18]. Similarly, in NC toxins, the NBP closure brings together ATP/CTP- and metal-ion-coordinating residues belonging to four distinct regions, which are distant from each other in the absence of cofactor and substrate. Three of these ATP/CTP- and metal-ion coordination regions are intrinsically flexible. Their positioning and stabilisation depend on both cofactor and substrate binding for switch A and B, and substrate binding for C_B_/LID (Figs 6A, 7A). Substrate-induced C_B_/LID closure after activation ultimately reduces the degrees of freedom of the nucleotide substrate and metal ions in the NBP (S7A-E Figs) to promote catalysis. Our mutational analysis in Vn-ExoY and the distantly-related Pa-ExoY identifies common elements of the NC toxin active sites that may contribute to catalysing the limiting steps in the cyclisation reaction with substrates of different sizes. The analysis pinpoints K3528^VnE-CA^/K81^PaE-CA^ and H3753^VnE-switch-B^/H291^PaE-switch-B^. Their mutations to Gln (to preserve polar interactions with ATP or Mg^2+^) strongly alter the AC/GC and CC/UC activity in Vn-/Pa-ExoY, respectively (Figs 6C-D, 7B-C). These residues are invariant in EF or CyaA and the same mutations on the equivalent K58^CyaA-CA^ and H298^CyaA-switch-B^ in CyaA [52, 53] or H577^EF-switch-B^ in EF [23] also largely reduce their CaM-stimulated AC activity. Our data show that the two residues are correctly positioned and stabilised with two metal ions only after cofactor and substrate binding and LID/C_B_ closure (Figs 6A, 7A, S7A-E). Further investigation is required to determine whether H3753^VnE-switch-B^ (or the equivalent histidine in other NC toxins) is critical for initiating 3’OH deprotonation via Mg^2+^_A_, and K3528^VnE-CA^ (or equivalent) for the phosphoryl transfer step in a manner analogous to R1029^tmAC-IIC2^ in tmAC. A key element in the ability of Vn-ExoY, and most likely other NC toxins, to catalyse the cyclisation reaction of nucleotides of different sizes is the tight coordination of the triphosphate moiety of purine and pyrimidine nucleotides after closure of their C_B_/LID subdomain. In the structure of Vn-ExoY bound to actin and 3’dCTP-2Mg^2+^, the hydrogen bond network around the ribose and base is weaker than for 3’dATP. This probably reduces the stability of pyrimidine nucleoside binding in the NBP of activated Vn-ExoY and may explain why Vn-ExoY remains less efficient as CC than AC. It will be interesting to compare these structural insights into the broad NC activity of NC toxins with the functional specificities of newly identified bacterial PycC pyrimidine cyclases, which appear to contain conserved catalytic elements of class III ACs/GCs [16].

Despite the lack of tertiary and quaternary structural homology, class II (NC toxins) and class III ACs may share some important determinants of catalysis. Upon activation, the collapse of their different ATP/metal-ion coordination regions towards the active site brings together and constrains all interactions with the various parts of the nucleotide substrate and the metal ions. This open-closed transition of their catalytic domains may also facilitate the conformational changes of the substrate predicted by computational studies during the cyclisation reaction [19, 31, 32, 40]. For the catalytic domain of cofactor-activated NC toxins, the substrate-induced fit mechanism is controlled by the C_B_/LID subdomain, which is near the cofactor-binding site. Its intrinsic flexibility and substrate-induced closure may ensure efficient enzymatic turnover through repeated closure and opening of the C_B_/LID for substrate loading/processing and product release (Fig 8). Without cofactor binding, substrate-induced NBP closure remains unproductive in catalysing the reaction because switch A and B, and thus part of the coordination of Mg^2+^ and the substrate nucleoside moiety (Figs 6A, 7A), are not properly stabilised. Our data suggest that the substrates enter the active site of NC toxins by binding through their triphosphate moiety. Indeed, the base-coordinating switch A is initially unstable and incorrectly positioned in ExoY toxins. This is consistent with the ability of NC toxins to use nucleotide substrates with different base sizes. Our analysis contributes to a deeper mechanistic understanding of how the functional dynamics of NC enzymes can be fine-tuned. Such high-resolution structural insights are still lacking for most NC enzymes, which are central components of numerous signalling cascades and relevant therapeutic targets for human pathologies involving cAMP/cGMP-signalling dysregulation [19, 21, 22, 54]. Our data reveal how the NBP of NC toxins gradually transitions to a catalytic conformation upon cofactor and substrate binding. These findings open up new structural perspectives for identifying ways to inhibit these NC toxins, which are involved in a variety of bacterial infections [1, 2, 6, 7].

## Materials and Methods

### Protein expression and purification

For biochemical studies, unless otherwise indicated, we used the 51.6-kDa Vn-ExoY functional module from V. nigripulchritudo MARTX (Vn-ExoY^3412-3873^) corresponding to its residues 3412-3873 (numbering from Uniprot accession no. (AC) F0V1C5) [5]. Vn-ExoY^3412-3873^ has a high sequence similarity (>85%) to all MARTX ExoY modules of *Vibrio* strains (Fig. S10). For Pa-ExoY, we used a highly soluble maltose-binding protein (MBP) fusion of the inactive mutant Pa-ExoY^K81M^ [3] in actin polymerisation assays. In enzymatic activity assays, we used Pa-ExoY constructs containing residues 20 to 378 from Unitprot AC O85345 with an N-terminal twinned-Strep-Tag (ST2). The fusion proteins of Pa-ExoY and Vn-ExoY with MBP or Trx were more suitable for dose-response analyses at high concentrations. They were more soluble than the isolated ExoY modules.

All Vn-/Pa-ExoY constructs (wild-type and mutants) were similarly expressed and purified under non-denaturing conditions using their Strep-Tag as previously described [3, 5]. The construct of Vn-ExoY (wild-type) or Vn-ExoY-K3528M-K3535M (Vn-ExoY^DM^) with an N-terminal Thioredoxin (Trx) (designated Trx-Vn-ExoY/Trx-Vn-ExoY^DM^, respectively) was customised as previously described with the N-terminal MBP fusion of Vn-ExoY [5]. The fusion constructs of Vn-ExoY and Pa-ExoY (Pa-ExoY residues 20 to 378 from Unitprot AC O85345) with an N-terminal twinned-Strep-Tag (ST2) or Thioredoxin (Trx, 11.6 kDa), designed as follows: ST2-(PreScission-cleavage-site)-(Vn-/Pa-ExoY) (denoted Vn-/Pa-ExoY in the text and cloned by Genscript) or HisTag-Trx-(PreScission-cleavage-site)-(Vn-ExoY)-ST2 (denoted Trx-Vn-ExoY) were cloned into pET29b vector using BamHI and XhoI restriction sites [5]. Expression was carried out in BL21 (DE3) bacteria grown in Luria-Bertani (LB) at 37 °C with shaking until an optical density at 600 nm of 0.6 was reached. After a heat shock on ice up to 16 °C, expression was induced by 1 mM β-isopropylthio-D-galactoside (IPTG) for 15 h (overnight) at the post-heat shock temperature, i.e. 16 °C. Cells were washed in PBS buffer, centrifuged, frozen in liquid nitrogen and stored at -20 °C until purification. Purified Vn-ExoY constructs were stored in 15 mM Tris pH 8.0, 100 mM KCl, 2 mM MgCl_2_, 1 mM DTT and purified Pa-ExoY constructs were stored in 20 mM Tris pH 8.5, 400 mM NaCl, 5 mM MgCl_2_, 1 mM DTT. The ST2- or Trx-fusion constructs of Vn-ExoY and Pa-ExoY showed similar specific activities for cAMP and cGMP synthesis, respectively, as the PreScission protease-cleaved Vn- or Pa-ExoY constructs. The wild-type and mutant Vn-/Pa-ExoY proteins have identical circular dichroism spectra and elute as monomeric proteins on calibrated size-exclusion chromatography columns.

α-actin (UniProt P68135) was purified from rabbit skeletal muscle as previously described [3, 55] and stored in G-buffer (5 mM Tris-HCl, pH 7.8, 1 mM DTT, 0.2 mM ATP, 0.1 mM CaCl_2_, and 0.01 % NaN_3_). Actin was pyrenyl-labelled at cysteine 374 [56]. Human profilin [57], CP [58], mDia1 FH1-FH2 [59], human VASP [60], Thymosin β4 [36], WH2 domain, and full-length human gelsolin [57] were used. Spectrin-actin seeds were purified from human erythrocytes as previously described [61]. ADP-actin was prepared from Ca-ATP-actin and converted to Mg-ADP-actin as previously described [3]. Latrunculin A was purchased from tebu-bio (produced by Focus Biomolecules). A 2-fold excess was added to G-actin and incubated for 10 min at room temperature before use in the activity assays.

### Pyrene-actin polymerisation assays

Actin polymerisation was monitored at 25 °C by the increase in fluorescence of 2-10 % pyrenyl-labelled actin (λexc = 340 nm, λem = 407 nm) using a Safas Xenius model FLX spectrophotometer (Safas, Monaco) with a multiple sampler device. Actin-Ca-ATP was converted to G-actin-Mg-ATP by the addition of 1/100 (vol./vol.) of 2 mM MgCl_2_ and 20 mM EGTA just prior to the experiments. Spontaneous ATP-G-actin polymerisation assays contained 4 µM G-actin (5 % pyrene-labelled) and were performed in a final buffer containing 50 mM KCl, 2 mM MgCl_2,_ 6 mM Tris-HCl pH 8, 1 mM TCEP, 1.5 mM ATP, 0.5 mM cAMP and additionally 0.5 mM GTP for Pa-ExoY experiments. Spontaneous ADP-G-actin polymerisation reactions containing 10 µM G-actin (10 % pyrene-labelled) were performed in a final buffer containing 30 mM KCl, 2 mM MgCl_2_, 5 mM Tris-HCl pH 7.8, 0.5mM cAMP, 3mM TCEP. Seed polymerisation reactions contained 1–2 μM actin (3-10 % pyrene-labelled), spectrin- or gelsolin-seeds, profilin, CP, VASP, formin mdia1 as indicated and were performed in a buffer containing 15 mM Tris pH 7.8 (or HEPES pH 7.5), 50-100 mM KCl, 2-4.5 mM MgCl2, 1-1.5 mM ATP, 1 mM cAMP and 2 mM TCEP.

The initial rates of filament growth from the barbed or the pointed ends were measured using spectrin-actin seeds for barbed end elongation or gelsolin-actin seeds for pointed end elongation. They were determined from linear fits to data collected during the first 2-3 min of elongation. These reactions contained 1.5 µM G-actin (5 % pyrene-labelled) and were performed in a final buffer containing 28 mM KCl, 2 mM MgCl_2_, 10 mM Tris-HCl pH 7.5, 2 mM ATP, 1 mM DTT.

Purely sequestering proteins form a CA complex with G-actin that do not participate in either barbed or pointed end growth. The following equation describes the changes in the initial growth rate, denoted v.

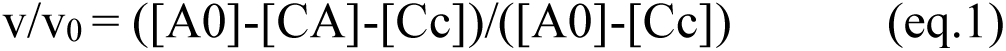

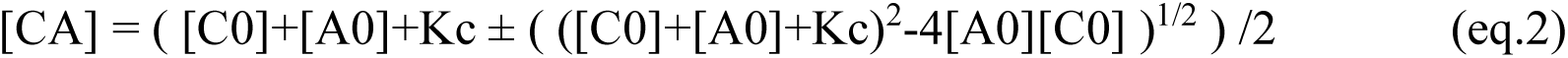

where [Cc] is the critical concentration, [A0] is the total concentration of G-actin in the growth assay and [CA] is the concentration of complex, [C0] is the total concentration of peptide, and Kc is the equilibrium dissociation constant between the protein (C corresponding to Vn-ExoY) and G-actin (A). V0 is the growth rate measured in the absence of protein C (V0=k_+_[F]([A0]-Cc), where k_+_ is the rate constant for G-actin association with barbed-ends and [F] is the concentration of F-actin seeds).

Kc was derived by fitting the dependence of the growth in eq1 on C concentration. Vn-ExoY leads to complete inhibition of pointed and barbed end growth by binding to G-actin with similar affinity in barbed and pointed end growth assays (Fig. 1D).

The effect of the chimera VnE^DM^-PRM-Prof on the initial rate of barbed end elongation of pre-assembled actin-spectrin seeds (0.3 nM) was determined by a linear fit to the first 150 seconds of the reaction performed by using 2 µM G-actin (3 % pyrene-labelled) in a final buffer containing 50 mM KCl, 2 mM MgCl_2,_ 50 mM Tris-HCl pH 7.8, 0.5 mM cAMP, 1 mM ATP and 2 mM TCEP (Fig. 3C).

The effect of Vn-ExoY on the initial rate of barbed end elongation of pre-assembled actin-spectrin seeds (0.36 nM) with increasing concentrations of profilin in the absence and presence of Vn-ExoY^wt^ at the indicated concentrations was determined by a linear fit to the first 150 s of assembly (Fig. 2D). These reactions were performed using 2 µM G-actin (5 % pyrene-labelled) in a final buffer containing 100 mM KCl, 2 mM MgCl_2,_ 15 mM Tris-HCl pH 7.8, 1 mM cAMP, 1 mM ATP, and 2mM TCEP. Elongation rates were normalised to the barbed-end elongation rate in the absence of both profilin and Vn-ExoY. The plot of the effect of profilin on barbed end elongation was fitted using the equation:

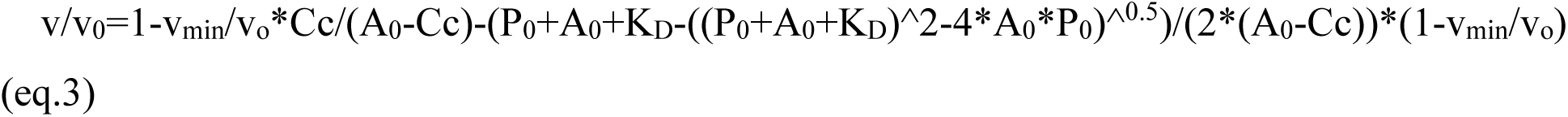

where v is initial elongation rate measured at profilin concentration [P], V_0_ initial elongation rate in the absence of profilin (V_0_=k_+_[F]([A_0_]-Cc), where k_+_ is the rate constant for G-actin association with barbed-ends and [F] is F-actin seed concentration), V_min_ initial elongation rate in the presence of saturating amounts of profilin, Cc critical concentration of barbed ends, A_0_ total G-actin concentration, P_0_ total profilin concentration and K_D_ profilin:G-actin equilibrium dissociation constant. In the presence of Vn-ExoY, good fits to the data were obtained by considering in (eq.3) that Vn-ExoY initially sequesters G-actin (preventing its assembly onto F-actin barbed ends) with or without profilin with a Kd of ∼12-14 µM as determined in Fig. 1d-e. This led to the use of the following equations:

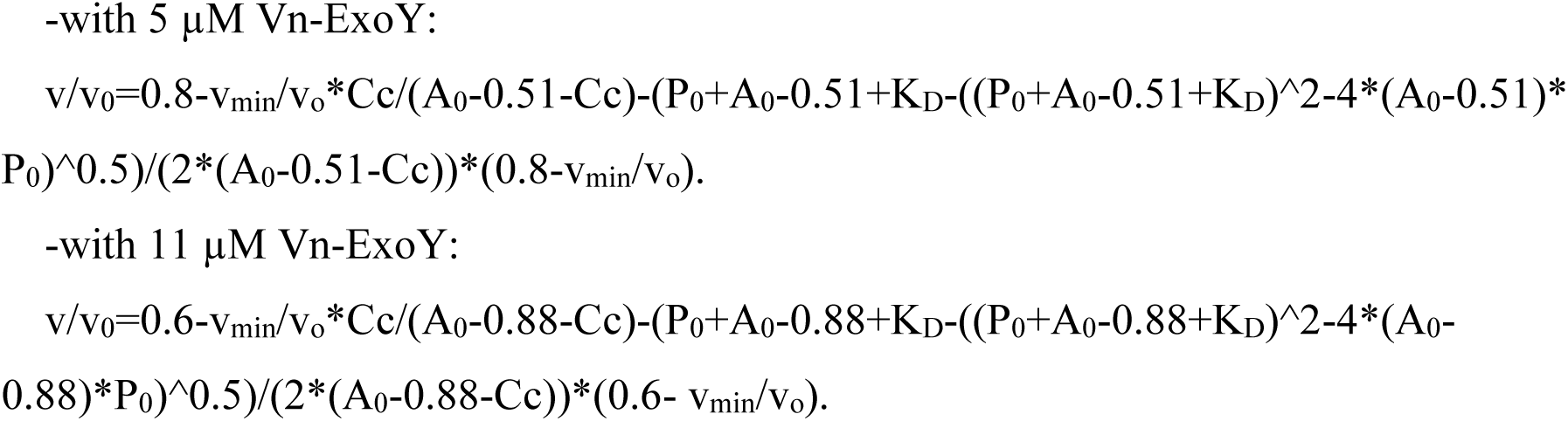

Polymerisation assays with mDia1 FH1-FH2 formin (Fig. 2E) and human VASP (Fig. 2F) were performed using 1 µM G-actin (5 % pyrene-labelled), profilin (8 µM), spectrin-seeds (0.13nM) in a final buffer containing 50 mM KCl, 4.7 mM MgCl_2_, 15 mM HEPES pH 7.5, 2.3 mM TCEP and 1.5 mM ATP. In experiments with VASP or formin in the presence of CP (2.4 nM), VASP (0.22 µM) or formin (3.2 nM), proteins were incubated in the reaction mixture for 2 min before the addition of CP, which was then incubated for a further 2 min. After the addition of CP, Vn-ExoY was added and fluorescence was recorded after 2 min incubation.

### Analytical Ultracentrifugation

Fluorescence-detected sedimentation velocity (FD-SV) experiments were performed using a Beckman XL-I (Beckman-Coulter, Palo Alto, USA) analytical ultracentrifuge (AUC) with an An-50Ti rotor equipped with an Aviv fluorescence detection system (AU-FDS, Aviv Biomedical). Actin was fluorescently labelled with Alexa-488 N-hydroxysuccinimide (NHS) ester (A20000, Thermo Fischer Scientific) as previously described [5].

Trx-Vn-ExoY, which resulted in higher purification yields and protein solubility at high concentrations, was preferred for titration FD-SV experiments. The buffer used for AUC experiments was (53 or 100 mM KCl, 15 mM Tris pH 7.5, 3 mM ATP, 4 mM CaCl_2_, 2 mM MgCl_2_, 0.5 mM Na-pyrophosphate, 2 µM latrunculin A, 1 % glycerol, 0.25 mg/mL BSA). A similar sedimentation coefficient (S_20,w_) was obtained for G-actin in both buffers. Samples and buffers were centrifuged at 15,000 g for 10 min prior to AUC experiments and solvent property measurements. Alexa-488 labelled G-actin at a final concentration of 60 nM (0.3 μM total G-actin, 20 % Alexa-488-labelled) was loaded into two-sector 12 mm path-length Epon charcoal-filled cells. 400 µL of samples were centrifuged at 40,000 rpm (116,369 g) at 20°C. Fluorescence data were collected at an excitation wavelength of 488 nm and emission wavelengths between 505 and 565 nm, with all cells scanned simultaneously every 6 min. Sedimentation velocity data were analysed using SEDFIT software [62] and the partial specific volume used for Alexa-488-G-actin was 0.723 mL/g. Buffer viscosities and densities were measured experimentally with an Anton Paar microviscosimeter and density meter (DMA 4500). Sedimentation velocity distributions were superimposed using GUSSI software [63] and peaks were integrated to obtain weighted-average binding isotherms (sw). The generated binding isotherms were then fitted using Sedphat software [64].

### Quantification of cAMP or cGMP synthesis in vitro

Vn- and Pa-ExoY-catalysed synthesis of cAMP, cGMP, cCMP or cUMP was measured in 50 µL reactions, as described previously [3, 5], containing 50 mM Tris pH 8.0, 0.5 mg/mL BSA, 200 mM NaCl, 1 mM DTT, MgCl_2_, 2 mM NTP (NTP corresponding to either ATP, GTP, CTP, or UTP) spiked with 0.1 µCi of [α-^33^P] NTP, Vn- and Pa-ExoY and indicated amounts of Mg-Actin-ATP.

### Microscale Thermophoresis experiments

Binding assays were performed in MST-optimised buffer equivalent to a low-ionic, non-polymerising G-buffer (5 mM Tris, pH 7.8, 0.1 mM CaCl_2_, 0.2 mM ATP) with 0.05 % Tween-20 on a Monolith NT.115 Microscale Thermophoresis (MST) device using standard treated capillaries (NanoTemper Technologies). Actin (final labelled-actin concentrations adjusted to 60 nM) was titrated from 108 to 0.033 µM by a 16-step 2-fold dilution series of Trx-Vn-ExoY^DM^. All binding experiments were repeated three times.

### Chimeric protein design containing Vn-ExoY, a short proline-rich motif (PRM), and profilin (VnE-PRM-Prof) for structural studies

We failed to crystallise our first functional Vn-ExoY^3412-3873^ construct [5] with ATP/ADP-G-actin, possibly because our functional MARTX Vn-ExoY fragment contained too many flexible regions to crystallise. To improve the crystallisation strategy, we designed a multi-modular chimeric protein containing C-terminally extended Vn-ExoY (residues 3455 to 3896), a short PRM/proline-rich motif capable of binding to profilin (consisting of five consecutive prolines with the C-terminal Pro3896 of Vn-ExoY and four additional prolines, and two glycines as a very short linker/spacer to connect the PRM to the N-terminus of profilin) and profilin (VnE-PRM-Prof in Figure 3A). The chimera VnE-PRM-Prof with an N-terminal twinned-Strep-Tag (ST2) and PreScission-cleavage-site (ST2-(PreScission-cleavage-site)-VnE-PRM-Prof) was cloned by Genscript. The VnE-PRM-Prof chimera was designed to: i) more effectively prevent actin self-assembly during crystallisation by sequestering G-actin simultaneously through Vn-ExoY and profilin; and ii) capture a functional four-component complex, some of which interact with only modest affinities. The chimeras were designed before the recent availability of the AlphaFold protein-prediction tool [65] or the recent cryo-EM structures of F-/G-actin-bound ExoY homologues [27]. Design was based on Pa-ExoY structure [25], ab initio models of Vn-ExoY, and protein-protein docking experiments with ab initio models of Vn-ExoY or Pa-ExoY and experimental distance constraints using FRODOCK software [66]. The distance constraints were based on the known contact regions between actin and Vn-ExoY or Pa-ExoY [3, 5] or structural similarities with CaM-activated EF and CyaA AC domains (e.g. to impose the proximity of switch A and C in the protein-protein interface) [23, 24]. Docking solutions needed to be filtered by experimental distance constraints to select a minimum number of equally likely solutions. We chose a long Vn-ExoY/profilin linker to accommodate different plausible docking solutions. We included a PRM to stabilise the Vn-ExoY C-terminus to profilin N-terminus junction and to study profilin-actin recruitment by PRMs of ABPs like formins or VASP. The chimera, named VnE-PRM-Prof, thus contains three main functional domains: Vn-ExoYwt/DM (residues Q3455 to P3896), a short PRM of 4 prolines (leading with Vn-ExoY P389 to the following sequence PPPPP (or P5), which allows profilin binding [67]) and human Profilin I sequence (Fig. 3A). The experiments shown in Fig. 3B and 3C confirmed the suitability of the chimera was for the analysis of Vn-ExoY interaction and activation by profilin:actin and for the crystallization of stable complexes. Using the VnE-PRM-Prof^wt/DM^ chimeras, we obtained the first high-resolution diffracting crystals of functional complexes with actin-ADP and further optimised and shortened the Vn-ExoY boundaries (Fig. 3A, residues 3455-3863) for subsequent structural studies.

Upon publication, the conditions for protein crystallisation will be available for the 6 complexes whose structures have been determined.

### Data collection and processing

The data were collected at 100 K on the PROXIMA-1 and PROXIMA-2 protein crystallography beamlines at the SOLEIL synchrotron (Saint Aubin, Université Paris-Saclay, France) using the *MXCuBE* application [68] and checked on the fly using the *XDSME* interface [69]. The X-ray energy was set to the Mn *K*-edge energy of 6.64 keV for the complex 5 (12.6 keV was used for the other crystals). All data sets used for structure solution and refinement were reprocessed using *autoPROC* [70] running *XDS* [71] for data indexing and integration, *POINTLESS* [72], and *AIMLESS* [73] for data reduction scaling and calculation of structure-factor amplitude and intensity statistics. For complexes 4 and 5, respective data sets from four and three different isomorphous crystals respectively were merged. Finally, *STARANISO* [74] was used to process and scale the anisotropy of the diffraction data, assuming a local I/σ(I) of 1.2. Thus, with the exception of crystal 3, the diffraction was extended to higher angles in the **a**\*/**c**\* plane, suggesting the inclusion of useable data beyond the spherical resolution.

### Structure determination, refinement and analysis

The structures were solved by the molecular-replacement method with Phaser [75], using the structure of free Pa-ExoY [25] and the profilin-actin structure (PDB 2pav) [76] as search models to place one (complexes 1 to 3) or two copies (complexes 4 and 5) per asymmetric unit. The first initial model was automatically built using Buccaneer [77] and completed using Coot [78]. Refinement, leaving 5 % of the reflections for cross-validation, was performed with BUSTER [79] using non-crystallographic constraints for complexes 4 and 5 and the ‘one TLS group per protein chain’ refinement. Structure quality was analysed using *MolProbity* [80]. The anomalous data set from complex 5 was analysed using ANODE [81]. Table S1 summarises the data collection, refinement statistics and the PDB accession codes. All figures showing protein structures (including the porcupine plot in Fig. 4A) were generated using *PyMOL* [82]. Motion analysis of protein subdomains was performed using DYNDOM [83].

The structure obtained with the shorter, optimised fragment of ExoY (residues 3455 to 3863) with the two proteins actin-ATP and profilin (Vn-ExoY-SO_4_^2-^:actin-ATP-LatB:profilin) is very similar to the structures obtained with the chimera VnE-PRM-Prof (VnE^DM^-PRM-Prof:actin-ADP-LatB or VnE-PRM-Prof-SO_4_^2-^:actin-ADP, Fig. 3A). It shows a similar disorder in the Vn-ExoY switch A, indicating that the switch A dynamics are not induced by the chimera.

The structural rearrangements in the Vn-ExoY switch regions observed between the Vn-ExoY^wt^-SO_4_^2-^:ATP-actin-LatB:profilin and Vn-ExoY^wt^-3’dATP-2Mg^2+^:ATP-actin-LatB structures are accompanied by minor changes elsewhere in the Vn-ExoY C_A_ subdomain, such as helices B (residues 3489-3504) and I (residues 3729-3742), which move ∼1.8 and ∼2.6 Å closer to switch A, respectively (Fig. 4A).

## Data Availability

Structural data have been deposited in the Protein Data Bank under PDB codes: 8BJH, 8BJI, 8BJJ, 8BR1, 8BO1, 8BR0. See S1 Table in the SI Appendix for further details.

## Acknowledgements

We thank SOLEIL for providing synchrotron radiation facilities, PROXIMA-1 and PROXIMA-2 beamline staff, and Pierre Legrand for strong technical support during data collection. We acknowledge the core facilities of Imagerie-Gif (http://www.i2bc.paris-saclay.fr).

## Author contributions

L.R. designed research. M.T.N, M.C., A.A.M., and L.R. performed the molecular biology or biochemical experiments, C.V. the AUC experiments, M.T.N, P.R., S.P., and L.R. the crystallisation and/or crystallographic experiments. D.R.B. and U.M. performed NC activity assays and generated the Pa-ExoY mutants. L.R. and M.T.N. designed the figures and wrote the paper. All authors reviewed and edited the article.

