## Supplementary material for "Functional and structural insights into the multi-step activation and catalytic mechanism of bacterial ExoY nucleotidyl cyclase toxins bound to actin-profilin": Supp. Table 1, Supp. Fig. S1 to S10

Short title: Mechanistic insights into ExoY NC toxin activation by actin-profilin

Magda Teixeira-Nunes[1], Pascal Retailleau[2], Dorothée Raoux-Barbot[3], Martine Comisso[1], Anani Amegan Missinou[1], Christophe Velours[1,#a], Stéphane Plancqueel[1], Daniel Ladant[3], Undine Mechold[3], Louis Renault[1*]

[1] Université Paris-Saclay, CEA, CNRS, Institute for Integrative Biology of the Cell (I2BC), 91198, Gif-sur-Yvette, France

[2] ICSN, CNRS, Université Paris Saclay, 1 avenue de la terrasse, 91190, Gif-sur-Yvette, France.

[3] Institut Pasteur, Université Paris Cité, CNRS UMR 3528, Unité de Biochimie des Interactions macromoléculaires, Département de Biologie Structurale et Chimie, 25-28 rue du Docteur Roux, 75724 Paris cedex 15, France

[#a] present address: Fundamental Microbiology and Pathogenicity Laboratory, UMR 5234 CNRS-University of Bordeaux, SFR TransBioMed, 33076 Bordeaux, France

This PDF file includes:

- Tables S1
- Legend for Movie S1
- Figures S1 to S10

| Complex | 1 | 2 | 3 | 4 | 5 | 6 |
|---|---|---|---|---|---|---|
| Complex | **Vn-ExoY (residue# Q3455 to P3896 with K3528M, K3535I) fused to PRM-profilin: ADP-actin-LatB** | **$SO_4^{2-}$-bound Vn-ExoY (residue Q3455 to P3896) fused to PRM-profilin: ADP-actin** | **$SO_4^{2-}$-bound Vn-ExoY (wt) (residue Q3455 to L3863): ATP-actin-LatB: profilin** | **(3'dATP-$2Mg^{2+}$)-bound Vn-ExoY (wt) (residue# Q3455 to L3863): ATP-actin-LatB** | **(3'dATP-$2(Mn/Mg)^{2+}$)-bound Vn-ExoY(wt) (residue Q3455 to L3863): ATP-actin-LatB** | **(3'dCTP-$2(Mn)^{2+}$)-bound Vn-ExoY(wt) (residue Q3455 to L3863): ATP-actin-LatB** |
| **PDB code** | **8BJH** | **8BJI** | **8BJJ** | **8BR1** | **8BO1** | **8BR0** |
| Synchrotron / beamline | SOLEIL / PROXIMA-1 | SOLEIL / PROXIMA-1 | SOLEIL / PROXIMA-2 | SOLEIL / PROXIMA-1 | SOLEIL / PROXIMA-2 | SOLEIL / PROXIMA-1 |
| Date of data collection | 12.06.2020 | 03.07.2020 | 06.12.2020 | 26.03.2021 | 26.03.2021 | 22.04.2022 |
| Wavelength (Å) | 0.979 | 0.979 | 0.980 | 0.979 | 1.893 | 0.979 |
| Resolution range (Å)in data reduction | 93.57–1.69 (1.85–1.69) | 61.81–1.64 (1.82–1.64) | 44.20–1.70 (1.73–1.70) | 47.76-2.04 (2.21-2.04) | 48.24-2.50 (2.70-2.50) | 48.05-2.22 (2.36-2.22) |
| Space group | $P2_1$ | $P2_1$ | $P2_1$ | $P2_1$ | $P2_1$ | $P2_1$ |
| *a*, *b*, *c* (Å) | 81.74, 62.82, 93.58 | 81.97, 62.85, 93.59 | 81.19, 63.20, 93.46 | 76.12, 132.44, 96.40 | 75.50, 132.17, 96.03 | 74.47, 132.50, 96.22 |
| α, β, γ (°) | 90, 91.03, 90 | 90, 90.28, 90 | 90, 91.04, 90 | 90, 110.57, 90 | 90, 110.80, 90 | 90, 110.40, 90 |
| Total reflections | 557546 (26053) | 584789 (29619) | 1584617 (71381) | 2829414 (138734) | 1287849 (68902) | 518884 (25532) |
| Unique reflections | 81375 (4069) | 83837 (4192) | 104207 (5211) | 86495 (4319) | 45059 (2239) | 62059 (2919) |
| Multiplicity | 6.9 (6.4) | 7.0 (7.1) | 15.2 (13.7) | 32.7 (32.1) | 28.6 (30.8) | 8.4 (8.7) |
| Completeness spherical (%) | 76.5 (16.3) | 72.2 (13.9) | 99.6 (95.3) | 76.6 (18.8) | 74.1 (18.0) | 71.8 (13.7) |
| Completeness ellipsoidal (%) | 94.9 (60.1) | 94.6 (65.1) | 99.8 (99.0) | 84.3 (32.1) | 93.7 (57.5) | 92.2 (49.5) |
| Mean $I/\sigma(I)$ | 11.2 (1.7) | 8.5 (1.6) | 25.3 (2.7) | 14.1 (1.4) | 16.2 (1.0) | 11.8 (1.5) |
| Wilson *B* factor ($Å^2$) | 26.5 | 22.6 | 24.1 | 51.1 | 75.8 | 64.2 |
| Matthews coefficient ($V_M$) ($Å^3Da^{-1}$) | 2.24 | 2.25 | 2.36 | 2.58 | 2.54 | 2.63 |
| Solvent content (%) | 45.18 | 45.36 | 47.88 | 52.35 | 51.61 | 53.18 |
| $R_{merge}$ (all I+ & I-) | 0.086 (0.910) | 0.123 (1.094) | 0.057 (1.055) | 0.162 (3.852) | 0.154 (4.021) | 0.098 (1.337) |
| $R_{meas}$ (all I+ & I-) | 0.093 (0.837) | 0.133 (1.182) | 0.059 (1.095) | 0.164 (3.914) | 0.157 (4.087) | 0.105 (1.420) |
| $R_{p.i.m.}$ (all I+ & I-) | 0.035 (0.384) | 0.051 (0.444) | 0.015 (0.289) | 0.029 (0.688) | 0.029 (0.734) | 0.036 (0.471) |
| $CC_{1/2}$ | 0.998 (0.725) | 0.996 (0.708) | 0.999 (0.86) | 0.999 (0.615) | 1.000 (0.575) | 0.998 (0.587) |
| Resolution range (Å) in refinement | 93.57–1.69 (1.80–1.69) | 25.06-1.75 (1.80-1.75) | 44.20–1.70 (1.71–1.70) | 47.76-2.04 (2.15-2.04) | 48.24-2.50 (2.63-2.50) | 48.05- 2.22 (2.36-2.22) |
| Reflections used in refinement | 81375 (1539) | 82105 (1573) | 104207 (1988) | 86494 (1654) | 45059 (863) | 62059 (1181) |

| Complex | 1 | 2 | 3 | 4 | 5 | 6 |
|---|---|---|---|---|---|---|
| Complex | **Vn-ExoY (residue[#] Q3455 to P3896 with K3528M, K3535I) fused to PRM-profilin: ADP-actin-LatB** | **$SO_4^{2-}$-bound Vn-ExoY (residue Q3455 to P3896) fused to PRM-profilin: ADP-actin** | **$SO_4^{2-}$-bound Vn-ExoY (wt) (residue Q3455 to L3863): ATP-actin-LatB: profilin** | **(3'dATP-$2Mg^{2+}$)-bound Vn-ExoY (wt) (residue[#] Q3455 to L3863): ATP-actin-LatB** | **(3'dATP-$2(Mn/Mg)^{2+}$)-bound Vn-ExoY(wt) (residue Q3455 to L3863): ATP-actin-LatB** | **(3'dCTP-$2(Mn)^{2+}$)-bound Vn-ExoY(wt) (residue Q3455 to L3863): ATP-actin-LatB** |
| **PDB code** | **8BJH** | **8BJI** | **8BJJ** | **8BR1** | **8BO1** | **8BR0** |
| Reflections used for $R_{free}$ | 4009 (89) | 4141 (70) | 5186 (97) | 4302 (76) | 2243 (39) | 2974 (61) |
| $R_{work}$ | 0.178 (0.221) | 0.175 (0.225) | 0.175 (0.225) | 0.190 (0.215) | 0.191 (0.235) | 0.208 (0.269) |
| $R_{free}$ | 0.212 (0.229) | 0.212 (0.269) | 0.195 (0.238) | 0.222 (0.240) | 0.231 (0.283) | 0.243 (0.375) |
| Total nb of atoms | 7980 | 7926 | 7763 | 12811 | 12368 | 12 638 |
| Macromolecules | 6900 | 6857 | 6922 | 11963 | 11927 | 11927 |
| Ligands | 55 | 33 | 64 | 182 | 182 | 170 |
| Solvent | 1025 | 1036 | 777 | 666 | 259 | 541 |
| No. of protein residues (per chain) | 366(A), 502(B) | 366 (A), 498 (B) | 367 (A), 355 (B), 139 (C) | 358, 365 (A, C), 398, 397 (B, D) | 360, 364 (A, C), 398, 396 (B, D) | 354, 365 (A, C), 398, 398 (B, D) |
| R.m.s.d., bond lengths (Å) | 0.008 | 0.008 | 0.008 | 0.008 | 0.008 | 0.008 |
| R.m.s.d., angles (°) | 0.92 | 0.92 | 0.96 | 0.93 | 0.98 | 0.97 |
| Ramachandran favored (%) | 96.4 | 96.4 | 96.6 | 97.9 | 95.8 | 96.5 |
| Ramachandran allowed (%) | 3.0 | 3.4 | 3.1 | 2.0 | 4.2 | 3.6 |
| Ramachandran outliers (%) | 0.6 | 0.2 | 0.3 | 0.1 | 0 | 0.1 |
| Average *B* factor (Å$^2$) | | | | | | |
| Overall | 35.0 | 30.3 | 33.4 | 61.7 | 85.1 | 72.0 |
| Macromolecules | *33.6* | *29.3* | *33.3* | *61.7* | 85.6 | 72.8 |
| Ligands | *24.3* | *23.5* | *25.2* | *51.6* | 77.1 | 56.0 |
| Solvent | *43.8* | *40.0* | *41.5* | *63.6* | 68.4 | 59.7 |
| No. of TLS groups | 2 | 1 | 3 | 4 | 4 | 4 |
| **PDB code** | **8BJH** | **8BJI** | **8BJJ** | **8BR1** | **8BO1** | **8BR0** |

(Values in parentheses are for the highest resolution shell).

[#]: First and last indicated residues in Vn-ExoY constructs are from the Uniprot A0A6N3LUE9_9VIBR sequence from *Vibrio nigripulchritudo* MARTX toxin.

**Table S1. Crystallographic data-collection and refinement statistics.**

**Movie S1. Rotation of the Vn-ExoY $C_B$/LID subdomain (red) towards its $C_A$ subdomain (black) upon 3’dATP binding between the nucleotide-free Vn-ExoY$^{wt}$-$SO_4^{2-}$actin-ATP-LatB:profilin and 3’dATP-bound Vn-ExoY-3’dATP-2*$Mg^{2+}$:actin-ATP-LatB structures.** **A)** Side view of both structures superimposed on their black $C_A$ subdomain. **B)** Frontal view. The axis of rotation (horizontal in (**A**)) is indicated by cyan spheres in a line. The $SO_4^{2-}$ ion and 3'dATP-2*$Mg^{2+}$ are represented by ball-and-stick models and their transparent molecular surfaces. Secondary structures are shown in yellow, except for those of switch A, B, and C, which are shown in blue, cyan, and purple, respectively. Actin:profilin and actin structures are omitted. Domain motion analysis performed with DYNDOM[1] identifies a 25° rotation of the $C_B$/LID subdomain of Vn-ExoY as a rigid body (in red, residues K3553-M3662$^{VnE\text{-}CB/LID}$) around the hinge regions 3532-3533$^{VnE}$ and 3662-3663$^{VnE}$ with respect to the $C_A$ subdomain (in black, residues 3468-3532$^{VnE\text{-}CA}$ and 3663-3849$^{VnE\text{-}CA}$ excluding the switch A, B, C, whose conformations vary between the two structures) (see also Fig. 4A,B).

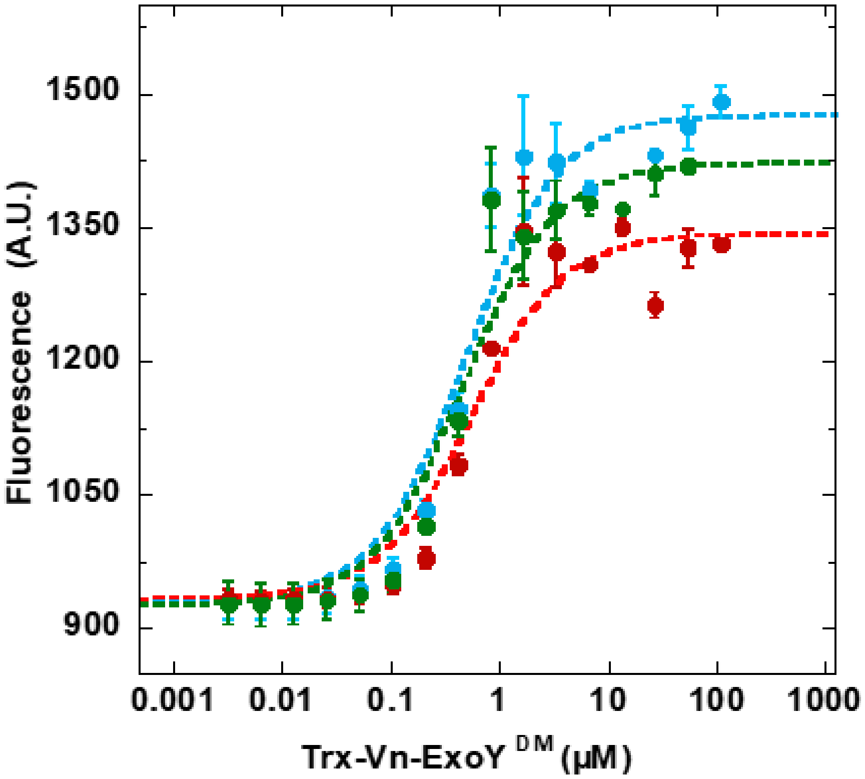

| Profilin (µM) | $K_d$ (µM) |
|---|---|
| 0 | 0.43 ± 0.11 |
| 4 | 0.51 ± 0.14 |
| 7.9 | 0.44 ± 0.11 |

**Fig. S1. Trx-Vn-ExoY$^{DM}$ binds free or profilin-bound G-actin with a very similar affinity.** Titration of free or profilin-bound Alexa-488-labelled ATP-G-actin with unlabelled Trx-Vn-ExoY$^{DM}$ and determination of Kd values from changes in microscale thermophoresis (MST) intensities. 0.075 µM Alexa-488-labelled, latrunculin-A-bound G-actin alone/free (cyan) or bound to 4 (red) and 7.9 (green) µM profilin was titrated with Trx-Vn-ExoY$^{DM}$ concentrations as indicated. This corresponds to a 2-fold dilution series from 108 to 0.033 µM with raw data in an insert. Error bars are s.d. (n≥3). The experiment using either the changes in MST (here) or fluorescence (Fig. 2C) intensity gave very similar Kd. The Kds averaged over both experiments (MST and fluorescence) were for Trx-Vn-ExoY$^{DM}$ binding to: free G-actin (cyan): Kd=0.46 ± 0.11 µM), G-actin bound to 4 µM profilin (red): Kd=0.42 ± 0.14 µM, and G-actin bound to 7.9 µM profilin (green): Kd=0.4 ± 0.11 µM.

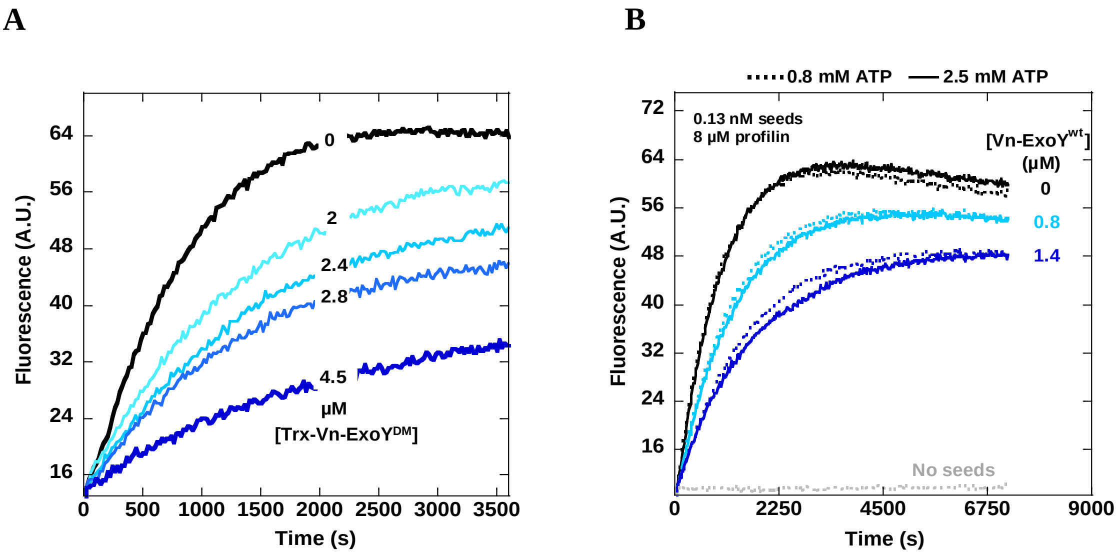

**Fig. S2. Vn-ExoY-induced inhibition of barbed-end growth in the presence of profilin-actin is independent of its adenylate cyclase (AC) activity. A)** Barbed end growth from spectrin-actin seeds (0.22 nM) was measured using 1 µM G-actin (5 % pyrenyl-labelled), 8 µM profilin, and the indicated concentrations (µM) of the inactive Trx-Vn-ExoY$^{DM}$ construct. Trx-Vn-ExoY$^{DM}$ inhibits barbed-end elongation from profilin-actin like Vn-ExoY$^{wt}$, demonstrating that Vn-ExoY-mediated effects on actin polymerisation are independent of its AC activity. The buffer was 50 mM KCl, 2 mM $MgCl_2$, 1 mM ATP, 15 mM Tris-HCl pH 7.8, 0.5 mM $CaCl_2$, and 1 mM TCEP. **B)** Barbed end growth from spectrin-actin seeds (0.13 nM) was measured using 1 µM G-actin (5 % pyrenyl-labelled), 8 µM profilin, and the indicated concentrations (µM) of active Vn-ExoY$^{wt}$ with either 0.8 (dashed lines) or 2.5 (solid lines) mM ATP. No differences in actin polymerisation kinetics were seen at different ATP concentrations. This indicates that the inhibitory effects of Vn-ExoY on actin polymerisation kinetics are independent of its AC activity throughout the duration of the experiments.

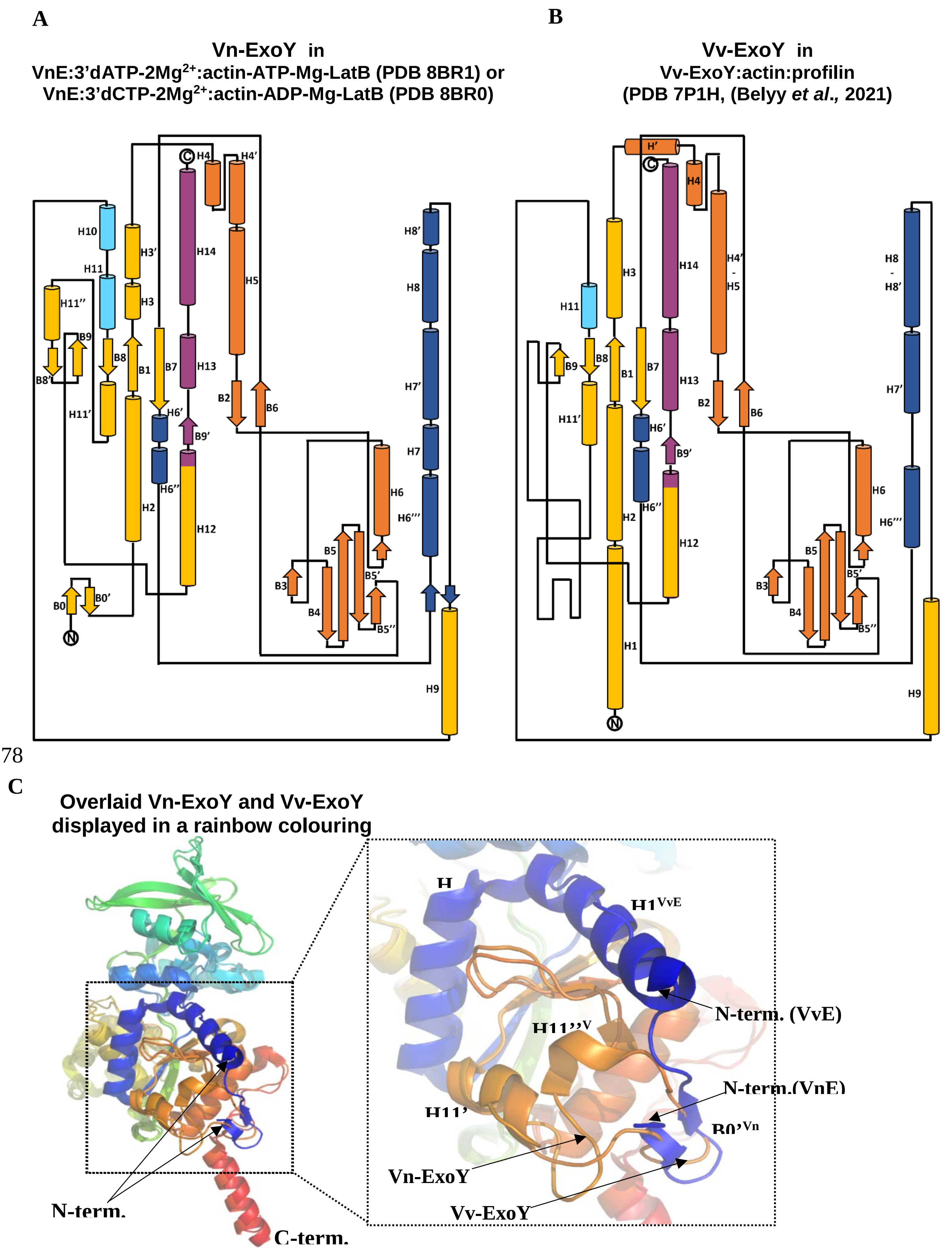
A
Vn-ExoY in
VnE:3'dATP-2Mg2+:actin-ATP-Mg-LatB (PDB 8BR1) or
VnE:3'dCTP-2Mg2+:actin-ADP-Mg-LatB (PDB 8BR0)
B
Vv-ExoY in
Vv-ExoY:actin:profilin
(PDB 7P1H, (Belyy et al., 2021)
H10
H11
H11''
B9
B8
B8'
H11'
H3'
H3
B1
H2
B7
H6'
H6''
H14
H13
B9'
H12
H4
H4'
H5
B2
B6
B0
B0'
B3
B4
B5
B5'
B5''
H6
H6'''
H8'
H8
H7'
H7
H9
H'
H4'-H5
H8-H8'
H1
C
Overlaid Vn-ExoY and Vv-ExoY
displayed in a rainbow colouring
N-term.
C-term.
H1VvE
N-term. (VvE)
N-term.(VnE)
H11''V
H11'
B0'Vn
Vn-ExoY
Vv-ExoY

**Fig. S3. Topology diagrams of (A) Vn-ExoY in PDB: 8BR1, 8BO1 or 8BR0, and (B) Vv-ExoY in PDB 7P1G [2].** (A) The structural topologies correspond in (A) to Vn-ExoY bound to actin-ATP-LatB and 3’dATP-2$Mg^{2+}$ (PDB 8BR1, 2.0 Å resolution), actin-ATP-LatB and 3’dATP-2$(Mn/Mg)^{2+}$ (PDB 8BO1, 2.5 Å resolution) or actin-ADP-LatB and 3’dCTP-2$Mn^{2+}$ (PDB 8BR0, 2.2 Å resolution), and in (B) to Vv-ExoY bound to actin-ATP-LatB and profilin (PDB 7P1G). In this 3.9-Å resolution cryo-EM structure, the 3'dATP ligand used for the preparation of the complex could not be modelled in Vv-ExoY. The amino-acid sequences of the Vn- and Vv-ExoY homologues are 89 % similar. However, their structures show a difference in the structural topology of the main chain trace at two segments of their sequence. (C) The superimposed structures of Vn-ExoY and Vv-ExoY, shown in a rainbow of colours (from blue to red from N- to C-terminus, respectively), with their N-terminus enlarged in the right inset. The backbone trace of Vn-ExoY N-terminal sequence $Y_{3468}$QSRDLVLE$P_{3477}$ overlaps with a small region located further in Vv-ExoY backbone. This overlapping region is located between switch B and C (namely between helix H11’ and β-stand B9) and corresponds to the Vv-ExoY sequence $L_{332}$GEGKGSIQ$T_{341}$. As a result, the backbone trace of their N-termini (Vn-ExoY sequence $K_{3466}$TYQSRDLVLEPIQHPKSIE$L_{3486}$, Vv-ExoY sequence $S_{19}$RDLVLEPIVQPETIE$L_{34}$) is very different, as is the backbone trace of their region between helix H11’ and β-stand B9 (Vn-ExoY sequence $D_{3781}$DGLGEGKGSIQ$T_{3793}$, Vv-ExoY sequence $E_{329}$DGLGEGKGSIQ$T_{341}$). The ab initio protein structure prediction of Vv-ExoY using the AlphaFold protein-prediction tool [3] suggests that Vv-ExoY adopts a backbone trace topology similar to that observed for Vn-ExoY in the crystal structures. Vv-ExoY bound to actin-ATP:profilin [2] and Vn-ExoY bound to actin-ATP/-ADP and 3’dATP/3’dCTP have otherwise similar overall conformations, including at their switch A, with a r.m.s.d. of 1.4 Å for 364 overlaid Cα atoms. The topology diagrams are adapted from those automatically generated by the Pro-origami [4] and PDBsum [5] programs.

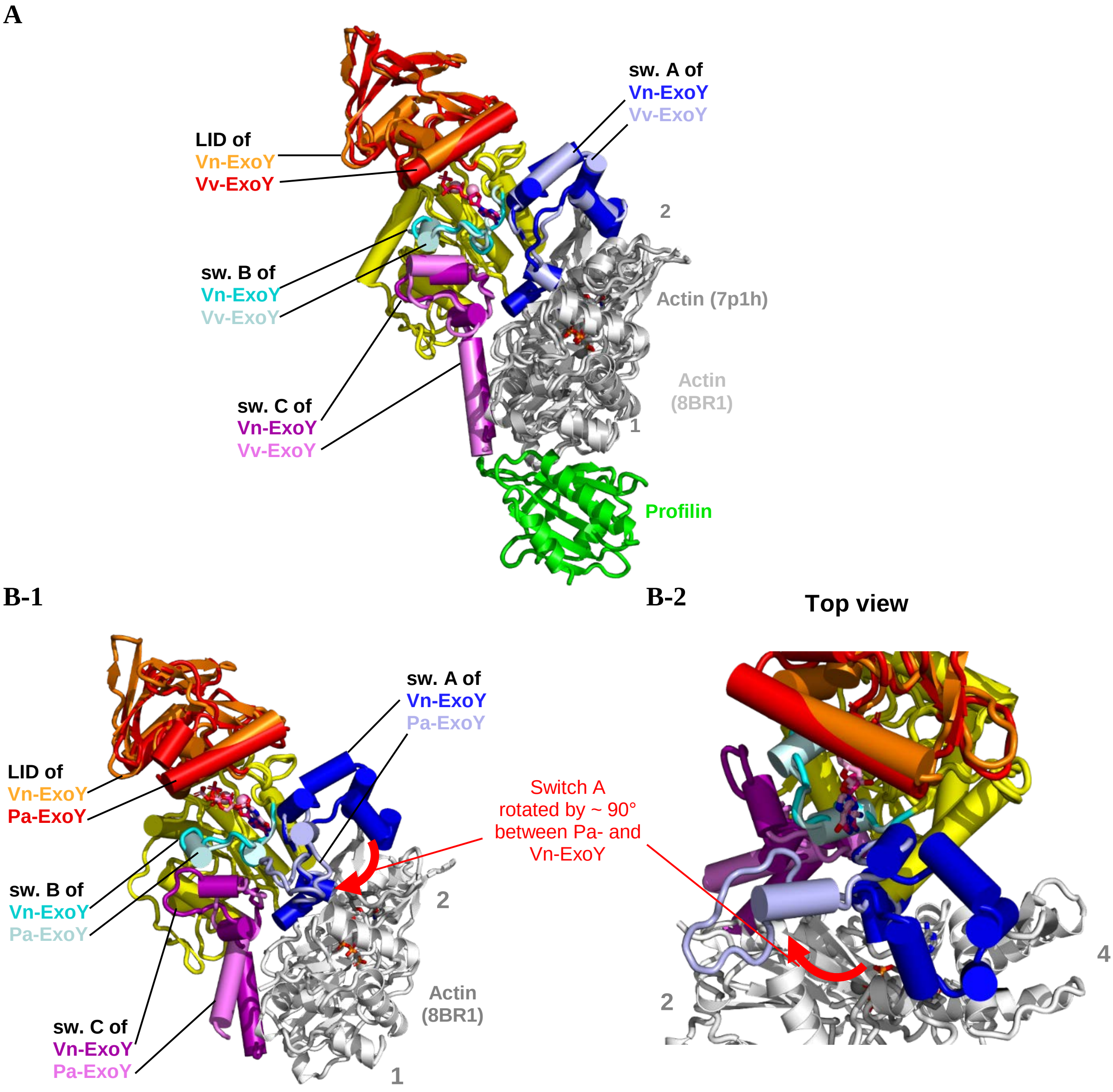

**Fig. S4. Overlays of the Vn-ExoY:3'dATP-2Mg$^{2+}$:actin-ATP-LatB structure (2.1 Å resolution crystal structure, PDB: 8BR1) with (A) the Vv-ExoY:actin-ATP:profilin structure (3.9 Å resolution cryoEM structure, PDB: 7P1H) [2] and (B) Pa-ExoY structure from the Pa-ExoY:3'dGTP-1Mg$^{2+}$:F-actin-ADP-Pi complex (3.2 Å resolution cryoEM structure, PDB: 7P1G) [2].** The structures are superimposed on their C$_A$ subdomain (residues 3468-3532$^{Vn\text{-}ExoY\text{-}CA}$ and 3663-3849$^{Vn\text{-}ExoY\text{-}CA}$). The four actin subdomains are indicated by grey numbers. Figures **(A)** and **(B-1)** show the same side views of the complexes, while **(B-2)** is a top view of **(B-1)**.

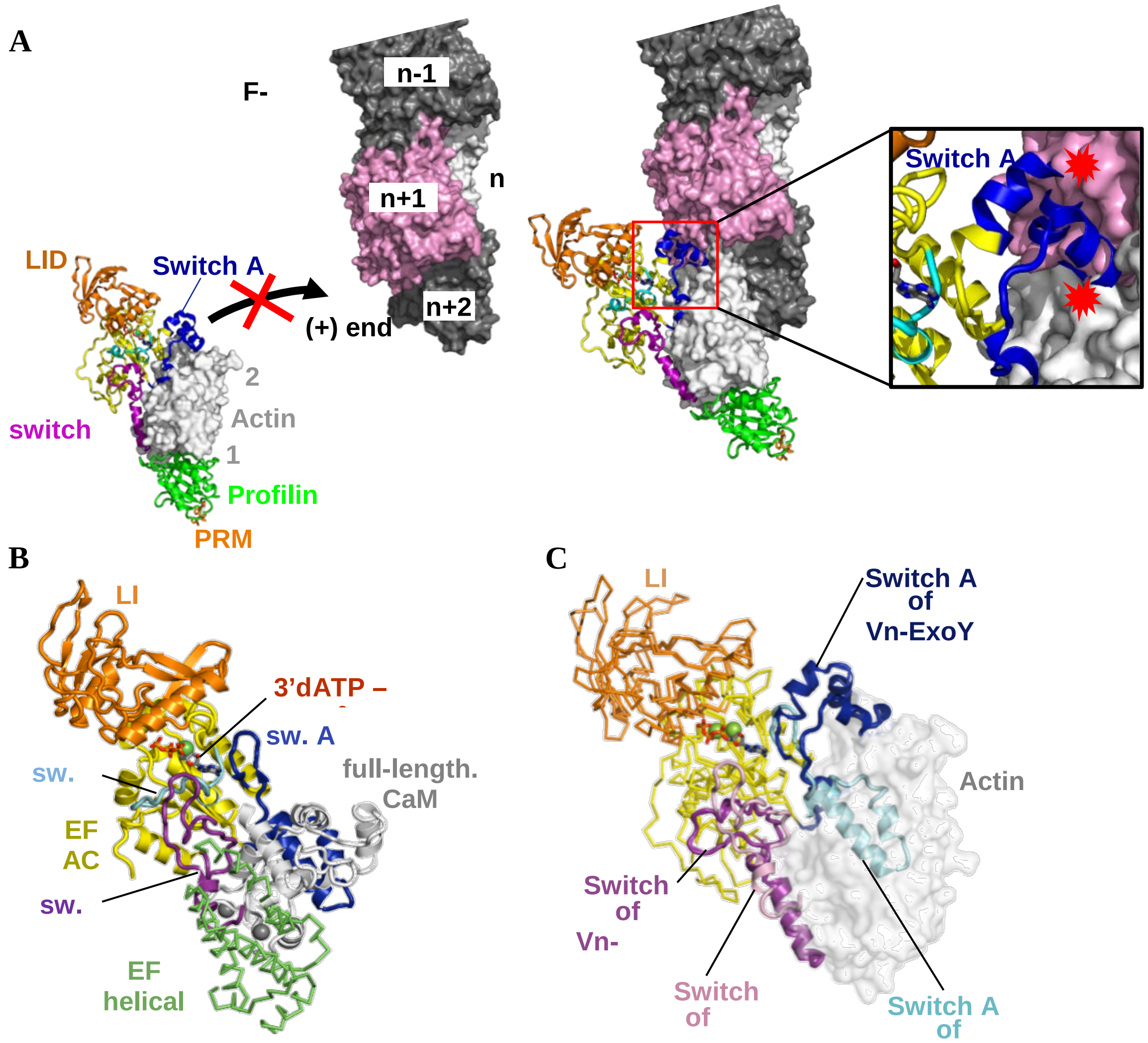

**Fig. S5. Switch A region is important for G-/F-actin and actin/calmodulin specificity. A)** Docking of the Vn-ExoY-3'dATP-2Mg$^{2+}$:actin-ATP structure in complex with profilin and PRM to F-actin (pdb 6FHL) shows that the interaction of the Vn-ExoY switch A between actin subdomains 2 and 4 prevents bound actin (in white) from assembling at the barbed (+) end of F-actin. It induces important steric clashes of switch A (red explosion symbols) with the penultimate actin subunit (in pink) at the barbed-end. Switch A binding to actin is also the region most incompatible with longitudinal contacts between adjacent actin subunits of the same F-actin strand (white and pink actin). This also most inhibits Vn-ExoY binding along F-actin [2, 6]. **B)** Cartoon representation of the 3'dATP-EF-CaM complex. The NC catalytic domains of EF, CyaA and Vn-ExoY use the same regions, i.e. switch A and C, and a similar orientation to interact with their cofactor. The C-terminal Ca$^{2+}$-binding globular domain of the 8.4-kDa protein CaM is responsible for most of the interactions with the EF or CyaA switch A and C [7-9]. However, it is much smaller than 42-kDa actin. It therefore only overlaps with actin subdomains 1 and 3. **C)** Structural comparison of G-actin- and CaM-activated Vn-ExoY and EF conformations, respectively. The common catalytic core, $C_A$ (yellow) and $C_B$ (orange)

domains are shown in a cartoon tube representation, and the regions determinant for cofactor specificity, switch A (pink and purple) and C (blue and cyan) regions, are shown in a cartoon representation. The positioning of switch A in EF or CyaA bound to CaM and in Vn-ExoY or Pa-ExoY bound to G or F-actin, respectively, is the most divergent region at the cofactor-toxin binding interface.

A

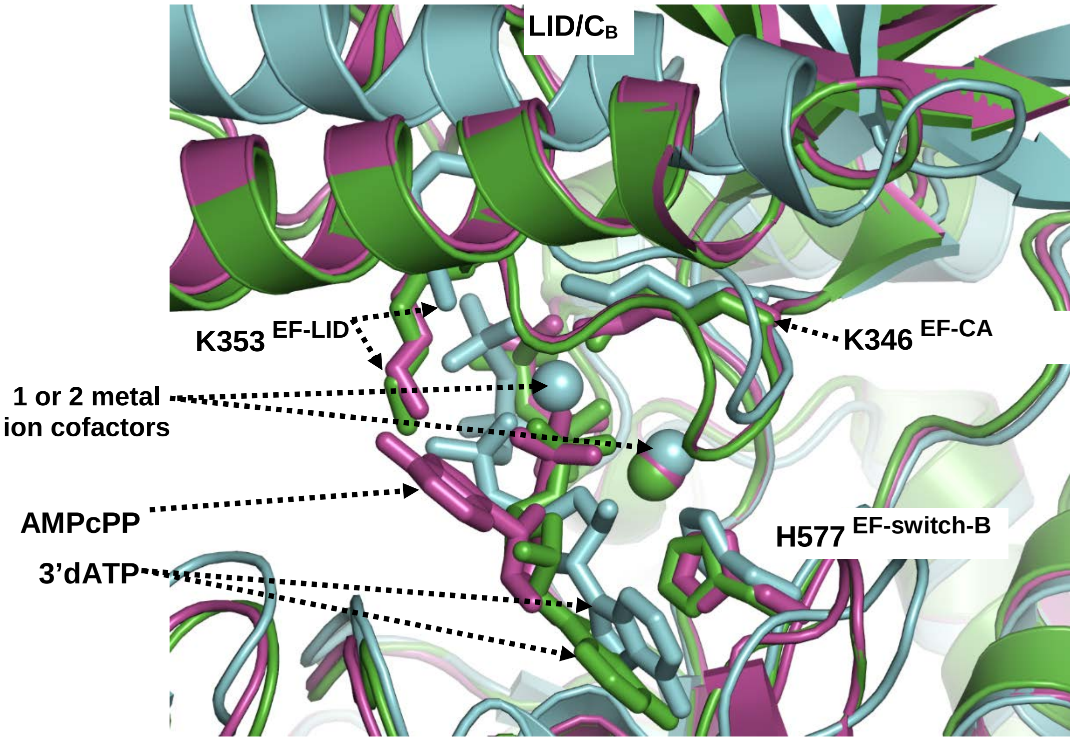

B

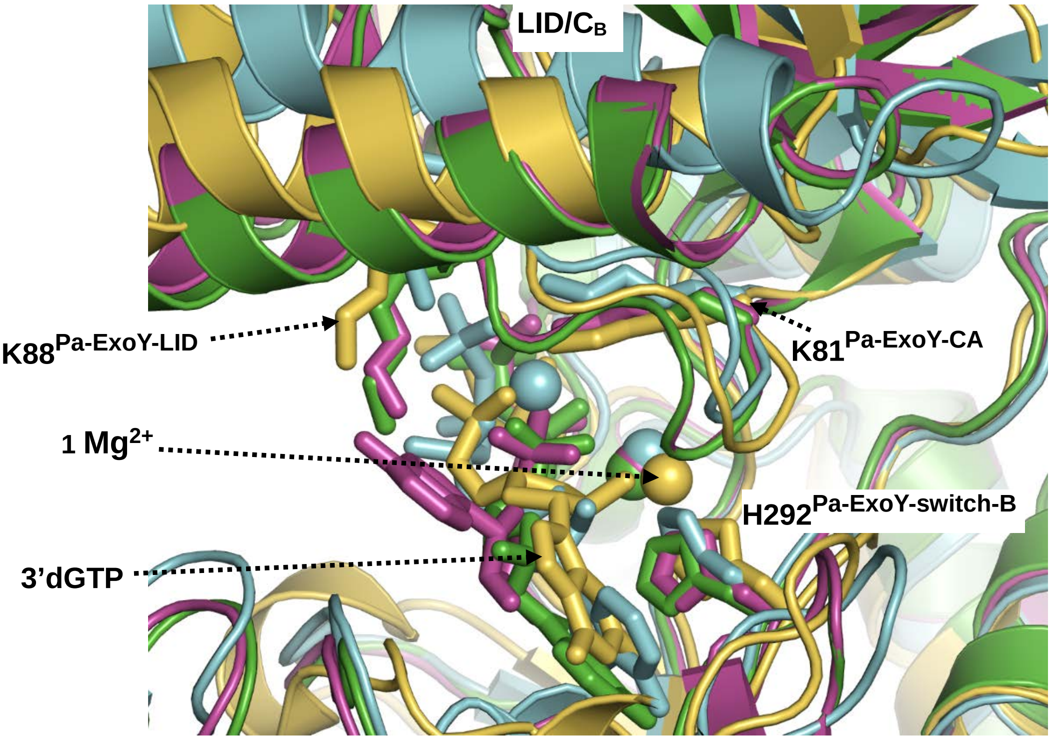

1XFV: CaM-activated EF: 3'dATP-2$Mg^{2+}$ LID open
1K90: CaM-activated EF: 3'dATP-1$Yb^{3+}$ LID closed
1S26: CaM-activated EF: AMPcPP-1$Yb^{3+}$ LID closed
7P1G: F-actin-activated Pa-ExoY: 3'dGTP-1$Mg^{2+}$ LID nearly-closed

**Fig. S6. The structures of EF bound to CaM and ATP analogue (A) or Pa-ExoY bound to F-actin and GTP analogue (B) show strong active-site structural heterogeneity.** They show either 1 or 2 metal ions associated with the substrate analogue, variable conformation or positioning of the different moieties of the purine nucleotide substrate analogues, and different positions of the LID/CB subdomain relative to the CA subdomain. The protein structures are shown in cartoon and coloured in cyan (pdb: 1XFV [9]), green (pdb: 1K90 [8]), light magenta (pdb: 1S26 [10]) and yellow (pdb: 7P1G [2]). The catalytic domains of the EF and Pa-ExoY NC toxins are superimposed on their $C_A$ subdomain. Metal ions are shown as spheres and the ligands and side chains of important catalytic residues ($K346^{EF\text{-}CA}$/$K81^{Pa\text{-}ExoY\text{-}CA}$, $H353^{EF\text{-}LID}$/$K88^{Pa\text{-}ExoY\text{-}LID}$ and $H577^{EF\text{-}Switch\text{-}B}$/$H292^{Pa\text{-}ExoY\text{-}Switch\text{-}B}$) are shown as sticks. **(A)** Overlay of the CaM-activated structures of EF bound to ATP analogues containing 1 or 2 metal cofactors. **(B)** Overlay of EF CaM-activated structures from (A) and Pa-ExoY F-actin-activated structure bound to 3’dGTP with 1 $Mg^{2+}$ metal ion.

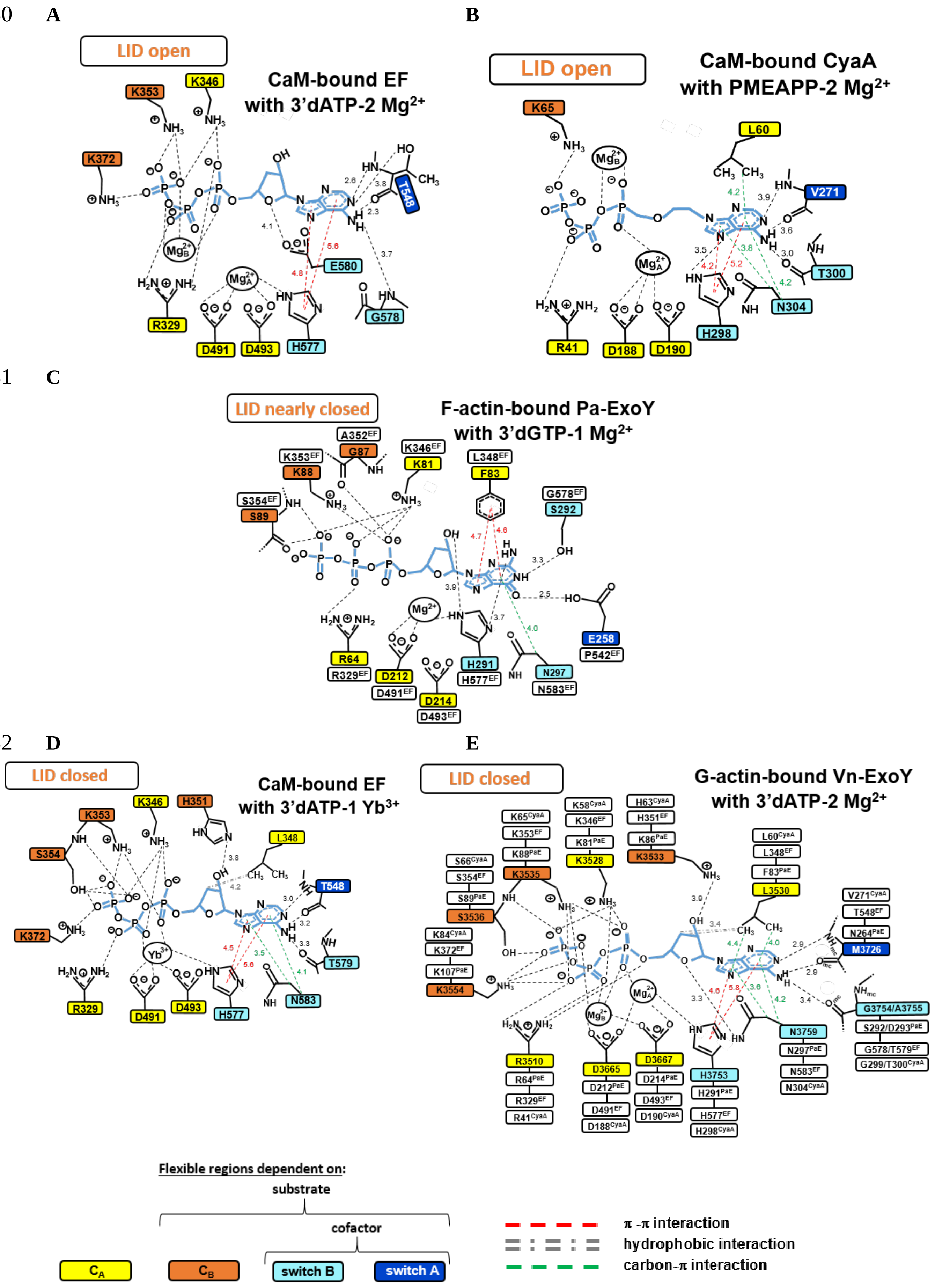
A
LID open
CaM-bound EF
with 3'dATP-2 Mg2+
B
LID open
CaM-bound CyaA
with PMEAPP-2 Mg2+
C
LID nearly closed
F-actin-bound Pa-ExoY
with 3'dGTP-1 Mg2+
D
LID closed
CaM-bound EF
with 3'dATP-1 Yb3+
E
LID closed
G-actin-bound Vn-ExoY
with 3'dATP-2 Mg2+
Flexible regions dependent on:
substrate
cofactor
switch B
switch A
π -π interaction
hydrophobic interaction
carbon-π interaction

**Fig. S7. Interactions with non-cyclisable ATP or GTP analogues and metal ions, and position of the LID/$C_B$ relative to the $C_A$ subdomain in CaM-bound EF/CyaA and F-actin-bound Pa-ExoY structures.** The active site of **(A)** CaM-activated EF with 3′dATP and two $Mg^{2+}$ metal ions (3.35 Å resolution crystal structure, PDB: 1XFV) [7], **(B)** CaM-activated CyaA structure with adefovir diphosphate (9-(2-(phosphonomethoxy)ethyl)adenine diphosphate, or PMEAPP) and two $Mg^{2+}$ metal ions (2.20 Å resolution crystal structure, PDB: 1ZOT) [9], **(C)** F-actin-activated Pa-ExoY with 3′dGTP and a single $Mg^{2+}$ metal ion (cryoEM structure at an average 3.20 Å resolution, PDB: 7P1G) [2], **(D)** CaM-activated EF structure with 3′dATP and a single $Yb^{3+}$ metal ion (2.75 Å resolution crystal structure, PDB: 1K90) [8], and **(E)** G-actin-activated Vn-ExoY with 3′dATP and two $Mg^{2+}$ metal ions (2.04 Å resolution crystal structure presented in this article, PDB: 8BR1). Thresholds for interaction detection are those of the PLIP (protein-ligand interaction profiler) [11] and Arpeggio [12] web servers, and those from the PoseView [13] tool available on the ProteinsPlus web server [14, 15]. Fig. S8 shows the position of the LID/$C_B$ relative to the $C_A$ subdomain in PDBs 1XFV (A), 1ZOT (B) and 1K90 (D). The use of an unconventional metal ion such as $Yb^{3+}$ for CaM-bound EF crystallisation is expected to alter the coordination of EF NBP with ATP only slightly compared to the putatitve physiological metal-ligand $Mg^{2+}$, as the metal $Yb^{3+}$ only slightly reduces the AC catalytic activity of EF [8].

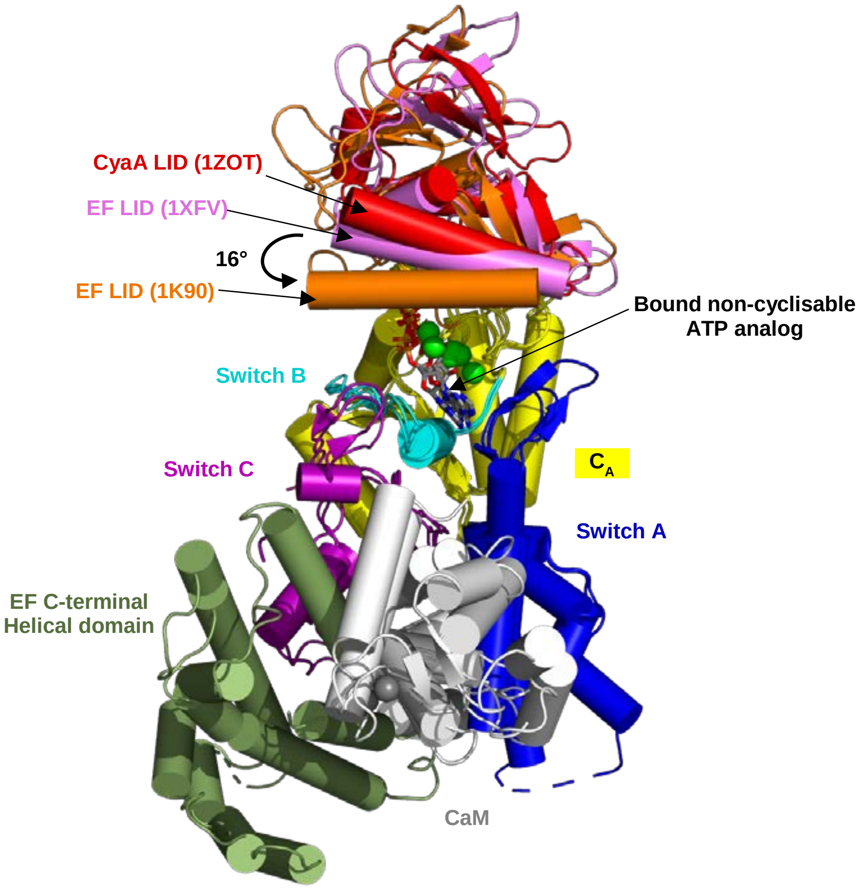

**Fig. S8. The LID/$C_B$ subdomain of CaM-activated EF and CyaA structures, in complex with various non-cyclisable ATP analogues, adopts different positions relative to the $C_A$ subdomain [7, 8], resulting in different positions and interactions with the ATP analogues and metal-ion(s) as shown in Figs. S6 and S7.** The EF structure with the LID in a closed conformation (1K90) and the EF and CyaA structures with the LID in an open conformation (1XFV and 1ZOT, respectively) are superimposed on their $C_A$ subdomains (in yellow). Protein domain motion analysis between the two structures of EF bound to CaM and 3'dATP (PDB 1K90 and 1XFV) shows that the LID/$C_B$ subdomain undergoes a 16° rotation as a rigid body around hinge regions $350\text{-}351^{EF}$ and $488\text{-}489^{EF}$ (motion analysis performed with DYNDOM [1]). Switch A, B, and C regions of EF and CyaA are coloured in blue, cyan, and purple, respectively. The LID region is shown in orange (1K90), pink (1XFV), and red (1ZOT). CaM and the additional C-terminal helical domain of EF are shown in white and green, respectively.

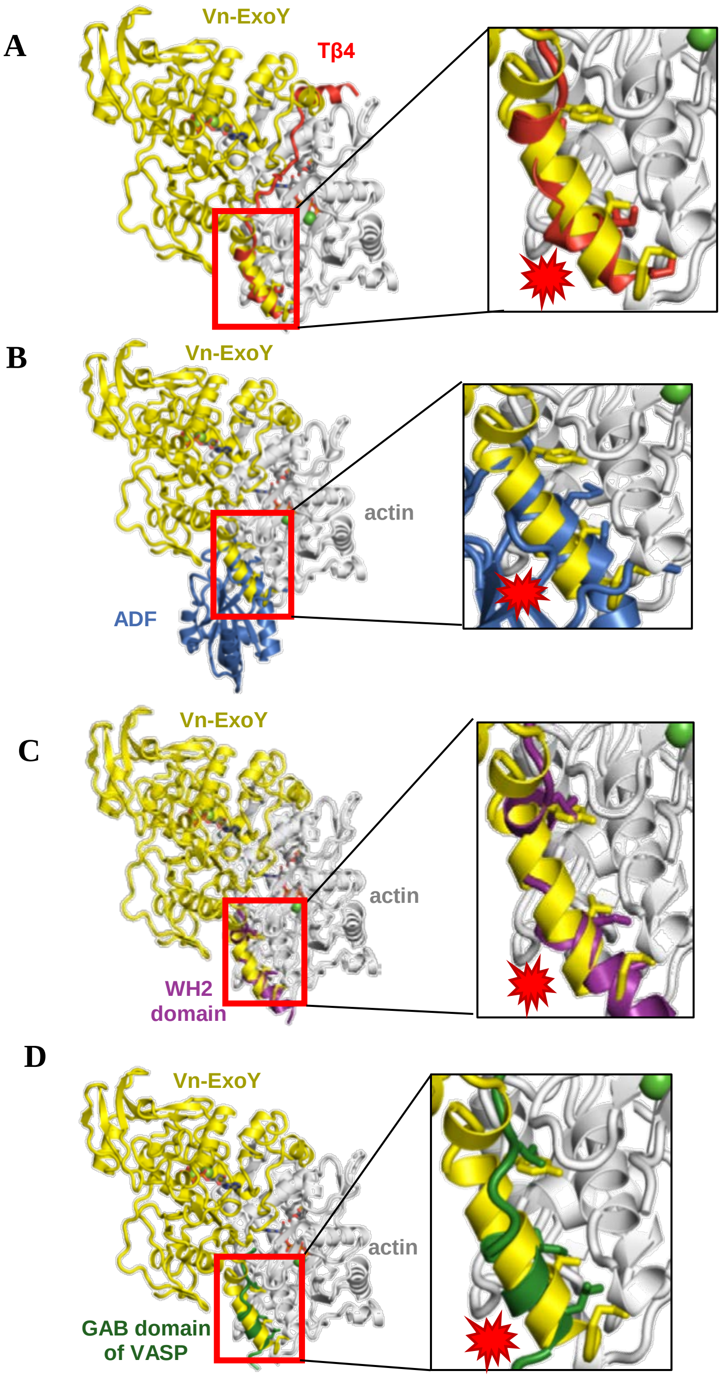

**Fig. S9. Binding interface overlap between Vn-ExoY switch C binding and binding of major G-actin-binding (GAB) proteins/domains: (A) Thymosin-β4 (Tβ4) (PDB: 4PL7 [16]), (B) ADF (PDB: 3DAW [17]), (C) Cordon-bleu WH2 domain (PDB: 5YPU [18]), (D) GAB domain of VASP (PDB: 2PBD [19]).** The interaction of the C-terminal amphipathic α-helix of Vn-ExoY switch-C within the hydrophobic cleft between actin subdomains 1 and 3 (shown in the right zoom panel) is a common binding site of actin-binding proteins (ABPs) [20]. Apart from profilin, this interaction of the Vn-ExoY switch-C C-terminus competes with a similar interaction that exists with most G-ABPs as shown in these examples. The structural models are consistent with Fig. 2A in the main text, which shows the competition in solution between Vn-ExoY and the Tβ4 or Cordon-bleu WH2 domain for binding to G-actin.

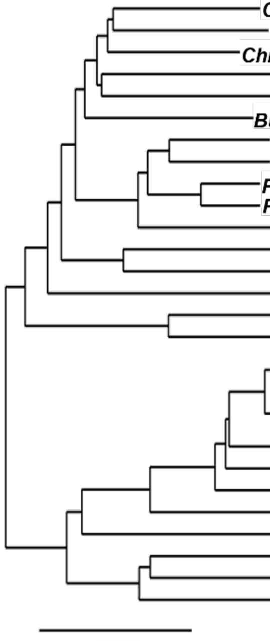

| Species | Accession | Sequence similarity with Pa-ExoY and Vn-ExoY (%) | Pa-ExoY: Solved PDB | Pa-ExoY: Switch A selectivity: G- / F-actin (G / F) | Vn-ExoY: Solved PDB | Vn-ExoY: Switch A selectivity: G- / F-actin (G / F) | Salt bridge with Asp25 Actin, Pa-ExoY-like: K347 | H352 | R364 | Vn-ExoY-like: R3824 | C-term. amphipathic α-helix | Catalytic residues: H291 PaE / H3753 VnE | K81 PaE / K3528 VnE |
|---|---|---|---|---|---|---|---|---|---|---|---|---|---|
| *Cedecea neteri* ExoY-L | WP_231562717 | 42 / 37 | | F | | | + | + | S | | + | + | + |
| *Yersinia aldovae* ExoY-L | WP_082167810 | 43 / 38 | | F | | | + | + | N | | + | + | + |
| *Chromobacterium vaccinii* ExoY-L | WP_115618495 | 42 / 38 | | F | | | + | + | + | | + | + | + |
| *Providencia stuartii* ExoY-L | WP_014658369 | 44 / 35 | | F | | | + | + | + | | + | + | + |
| *Salmonella enterica* ExoY-L | WP_079820003 | 44 / 39 | | F | | | R | Y | S | | + | + | + |
| *Burkholderia pseudomallei* ExoY-L | WP_154234830 | 43 / 38 | | F | | | + | + | + | | + | + | + |
| *Pseudomonas reactans* ExoY-L | WP_178953282 | 44 / 35 | | F | | | + | + | + | | + | + | + |
| *Pseudomonas viridiflava* ExoY-L | WP_122725039 | 42 / 36 | | F | | | + | + | + | | + | + | + |
| *Pseudomonas sivasensis* ExoY-L | WP_181639688 | 45 / 38 | | F | | | + | + | + | | + | + | + |
| *Pseudomonas grimontii* ExoY-L | WP_090407491 | 43 / 37 | | F | | | + | Y | + | | + | + | + |
| *Pseudomonas fluorescens* ExoY-L | WP_120450660 | 43 / 38 | | F | | | + | + | + | | + | + | + |
| *Pantoea cypripedii* ExoY-L | QGY31023 | 41 / 38 | | F | | | + | Y | S | | + | + | + |
| *Candidatus hamiltonella defensa* ExoY-L | ACQ67813 | 42 / 37 | | F | | | + | Y | S | | + | + | + |
| *Iodobacter fluviatilis* ExoY-L | QBC43103 | 39 / 35 | | F | | | + | + | + | | + | + | + |
| *Aeromonas salmonicida* ExoY-L | ALK43954 | 78 / 38 | | F | | | + | + | + | | + | + | + |
| *Pseudomonas aeruginosa* ExoY (Pa-ExoY) | WP_023129860 | 37 | (7P1G) | F | | | + | + | + | | + | + | + |
| *Vibrio anguillarum* ExoY-L | WP_020977741 | 88 / 37 | | | | G | | | | + | + | + | + |
| *Vibrio ordalii* ExoY-L | WP_069571922 | 89 / 37 | | | | G | | | | + | + | + | + |
| *Vibrio vulnificus* ExoY-L (Vv-ExoY) | WP_039507922 | 89 / 39 | | | (7P1H) | G | | | | + | + | + | + |
| *Vibrio cholerae* ExoY-L | WP_000517829 | 88 / 38 | | | | G | | | | + | + | + | + |
| *Vibrio nigripulchritudo* ExoY-L (Vn-ExoY) | WP_013610353 | 37 | | | (8BJJ, 8BR1)* | G | | | | + | + | + | + |
| *Vibrio neptunius* ExoY-L | WP_084720279 | 87 / 37 | | | | G | | | | + | + | + | + |
| *Psychromonas* sp.Urea-02u-13 ExoY-L | WP_101082559 | 87 / 37 | | | | G | | | | + | + | + | + |
| *Vibrio caribbeanicus* ExoY-L | EFP94682 | 70 / 36 | | | | G | | | | + | + | + | + |
| *Proteus vulgaris* ExoY-L | WP_232799119 | 59 / 37 | | | | G | | | | + | + | + | + |
| *Providencia alcalifaciens* ExoY-L | WP_159075030 | 55 / 35 | | | | G | | | | + | + | + | + |
| *Providencia rettgeri* ExoY-L | WP_105880947 | 53 / 37 | | | | G | | | | + | + | + | + |
| *Escherichia coli* ExoY-L | WP_153673629 | 53 / 35 | | | | G | | | | + | + | + | + |

**Fig. S10. Phylogenetic tree showing the relationships between ExoY-like proteins/effector domains produced by different β- and γ-proteobacteria.** ExoYs are classified as Vn-ExoY-like or Pa-ExoY-like homologues based on both their sequence similarity and predicted structure. Except for the indicated PDBs, the *ab initio* protein structure of all other ExoYs was predicted using the highly-accurate protein prediction tool AlphaFold [3]. The amino-acid sequence alignment of Pa-ExoY, Vn-ExoY and ExoY-like proteins/modules found in other γ- or β-proteobacteria and potentially activated by actin was performed using Clustal Omega (http://www.ebi.ac.uk, accessed on 12 September 2022) [21]. The multiple sequence alignment from Clustal Omega was used to construct the phylogenetic tree by using TreeDyn (www.phylogeny.fr) [22]. Pairwise sequence similarities (%) with Pa-ExoY and Vn-ExoY are shown in green and blue, respectively (Sequence Identity And Similarity (SIAS) tool, http://imed.med.ucm.es/Tools/sias.html accessed 12 September 2022). The NCBI accession numbers of the protein sequences are given to the right of the phylogenetic tree. The structures of Vn-ExoY solved in this study with actin or actin:profilin are marked with a red asterisk. The selectivity of the switch A conformation for G- or F-actin in the solved or AlphaFold-predicted structures of the ExoYs is indicated by G or F, respectively. The predicted structures of the ExoYs, which are closely-related to the MARTX Vn-ExoY module, are compatible with the formation of an ExoY:actin:profilin ternary complex and a conformation of switch A on actin that inhibits G-actin:profilin assembly at the barbed-ends of actin filaments. A + sign indicates that the Vn-ExoY or Pa-ExoY residue/structural element is conserved at the same position in the ExoY-like homologue's sequence and predicted structure.

### REFERENCES used in supplementary information:
